# Myopathy-causing variants of PYROXD1 converge on impaired RNA ligation

**DOI:** 10.64898/2026.10.09.758034

**Authors:** Sebastian Guzman-Perez, Moritz Kleinwächter, Muhammet Samet Özeren, Amelie Catharin, Stefan Weitzer, Milica Milašinović, Muhammed Buğra Yaman, Anastasiia Gorbenkova, Michaela Hahn, Rubina Koglgruber, David Hollenstein, Josef M. Penninger, Markus Hartl, Sandra T. Cooper, Frances J. Evesson, Javier Martinez, Igor Asanović

## Abstract

RNA ligation is critical for RNA processing pathways including pre-tRNA splicing. RTCB is the only essential RNA ligase in humans and the catalytic subunit of the tRNA ligase complex (tRNA-LC). Its active site is sensitive to oxidation and strictly depends on oxidoreductase PYROXD1, the only tRNA-LC subunit with multiple known pathogenic protein-coding variants. The role of RNA ligation in PYROXD1 myopathy remained contentious: loss of PYROXD1 primarily affects mitochondria, and ligation-linked readouts are inconsistent in existing assays. Here, we reveal that PYROXD1 is physically, genetically and functionally committed to the tRNA-LC. Selectively impairing RTCB activation reproduces hallmarks of PYROXD1 dysfunction, including mitochondrial deregulation. Using degron-based depletion of endogenous PYROXD1, we show that all six single amino acid variants reported in patients fail to sustain RNA ligation in proportion to their clinical severity. The variants segregate into three mechanistic classes: they lower protein stability, disrupt the NADPH-mediated conformational switch, or impair RTCB engagement by the C-terminal tail, a sequence unique to PYROXD1 and directly affected in the most severe variant D500N. Variant-level measurements in the degron model predict outcomes in homozygous and compound heterozygous patient fibroblasts. Together, these data establish impaired RNA ligation as the driver of PYROXD1 pathogenicity.

## Introduction

Ligation of RNA molecules entails head-to-tail joining of polyribonucleotide chains^1^. In humans, it occurs during pre-tRNA splicing^2^, the unfolded protein response (UPR)^3,4^, RNA repair^5^, and generation^6^ and suppression^7^ of transposons. The only essential metazoan RNA ligase is RTCB (HSPC117/TSLIG1)^8^, which recognises distinct RNA ends to facilitate selective RNA ligation, both for the endogenous substrates^9–11^ as well as in newly emerging transcriptome-editing technologies^12,13^.

Metazoan RTCB operates within the dynamic heterooligomeric tRNA ligase complex (tRNA-LC)^8,14^, consisting of three additional “core” proteins essential for complex stability (DDX1, CGI-99, and a FAM98 paralogue)^15,16^, the subcellular trafficking adaptor Ashwin^17^, and two functionally critical “expanded” complex subunits Archease^18^ and PYROXD1^19^. While Archease directly catalyses the initial step of the RNA ligation cycle^20^, PYROXD1 is required to keep the active site of RTCB in catalytically competent state including a reduced, metal-bound cysteine residue^21^.

Defects in RNA ligation were observed in multiple disorders, primarily affecting the nervous system. Such diseases are often triggered by variants of proteins more indirectly linked to the tRNA-LC such as CLP1^22–25^ or FMRP^26^, all of which have additional functions unrelated to RNA ligation. DDX1 harbours the single so far known published protein coding variant in a “core” tRNA-LC subunit, found in a patient displaying microcephaly^27^, and variants of an enhancer proximal to DDX1 gene show genomic association with orofacial clefts^28^. The indirect or non-specific involvement of the tRNA-LC, combined with lack of pathogenic variants in the “core” subunits, hampers the efforts to dissect the axis from a precise molecular defect to the clinical phenotype, and to specifically attribute them to insufficient RNA ligation.

In contrast, multiple reports have revealed that variants of the “expanded” subunit PYROXD1 cause a rare form of myopathy in humans^14,29–36^. Patients have either homozygous or compound heterozygous genotypes, with symptoms ranging from congenital debilitating loss of motor function to adult-onset moderate muscle weakness. PYROXD1 is hence the sole tRNA-LC component with multiple published pathogenic protein coding variants. Frameshift or exon skipping variants are presumably deleterious. However, the six single amino acid variants found in patients so far (see “variant nomenclature”) affect residues that are scattered across PYROXD1 structure, allowing for potential mechanistic dissection linking distinct molecular defects with severity of phenotypic outcomes. PYROXD1 could thus become the long missing key for understanding the molecular basis of diseases caused by defective RNA ligation.

The role of faulty RNA ligation in the pathology of PYROXD1 variants however remains contentious. RTCB only ligates nucleo-cytoplasmic RNAs^8,9,13,17^, while depletion of PYROXD1 appears to primarily affect mitochondria^30,32,33,36,37^. Some patient-derived fibroblasts displayed normal unfolded protein response^29^ and one pathogenic PYROXD1 variant was reported to normally interact with RTCB^21^. This questions whether the available readouts can fully assess the contribution of PYROXD1 variants to RNA ligation and if the loss of RNA ligation causes PYROXD1 myopathies.

Here, we demonstrate that PYROXD1 is largely bound to the tRNA-LC, and that the key cellular phenotypes upon its depletion, including mitochondrial deregulation, are mimicked by removal of the other tRNA-LC subunits. We establish the first system for comprehensive and background-free evaluation of cellular functionality of PYROXD1 variants, which revealed that all variants found in myopathy patients fail to fully sustain RNA ligation in a manner correlating with their pathogenicity. Variant failures fall into three mechanistic classes, linked with individual protein domains: FAD-binding domain variants decrease protein stability; NADPH-binding domain variants deregulate the nucleotide-mediated conformational switch; and C-terminal domain variants either kinetically delay RTCB association or abrogate the RTCB-binding interface. The most specific and severe variant D500N^31^ directly affects docking of the unique PYROXD1 C-terminal tail into the active site of RTCB. This thorough mechanistic dissection allowed us to successfully predict the effects of the variant G18R, so far only reported in a conference abstract^38^, establishing a solid groundwork for estimation and experimental testing of potentially pathogenic replacements in the future. Finally, segregation of PYROXD1 variants based on loss of RNA ligation allowed us to anticipate the defect in patient fibroblasts and establish a model that aligns ligation defects with disease onset. Together, these data establish the first comprehensive analysis of pathogenic variants in the tRNA-LC, revealing a shared cellular signature linking tRNA-LC dysfunction to mitochondrial proteomic alterations and indicating a direct and specific role of disrupted RNA ligation in PYROXD1-linked myopathies.

## Results

### PYROXD1 is physically and functionally committed to the tRNA-LC

To study the cellular consequences of depleting PYROXD1, we have previously employed RNA interference-based silencing in a cell line expressing short hairpin RNA (shRNA) against PYROXD1^19^. Near-total loss of PYROXD1 triggers a dramatic drop in double-stranded RNA oligonucleotide ligation in extracts, comparable to the loss of catalytic subunit RTCB and of Archease, the other “expanded” subunit involved in activation and protection of RTCB^18,20,21^ (**Fig. 1A**). Silencing of PYROXD1, RTCB and Archease also lowered efficacy of pre-tRNA fragment ligation (**Fig. S1A**), evident in accumulation of non-ligated fragments in the cells, without affecting cellular viability (**Fig. S1B**). In contrast to stress-induced mature tRNA fragmentation^39–41^, pre-tRNA fragments (also referred to as 5’ leader- and 3’ trailer-tRNA exons^22^) are constitutively generated as a part of pre-tRNA splicing^42–44^, a biogenesis process essential for maturation of intron-containing tRNAs. Unlike pre-tRNA splicing, *XBP1* mRNA splicing displayed higher data dispersion and is only mildly affected (**Fig. S1C**), in alignment with the previous observation that combined silencing of Archease and RTCB is required to substantially impact UPR^9^. This implies that monitoring *XBP1* splicing may not be suitable to reveal partial defects in RNA ligation.

**Figure 1.**
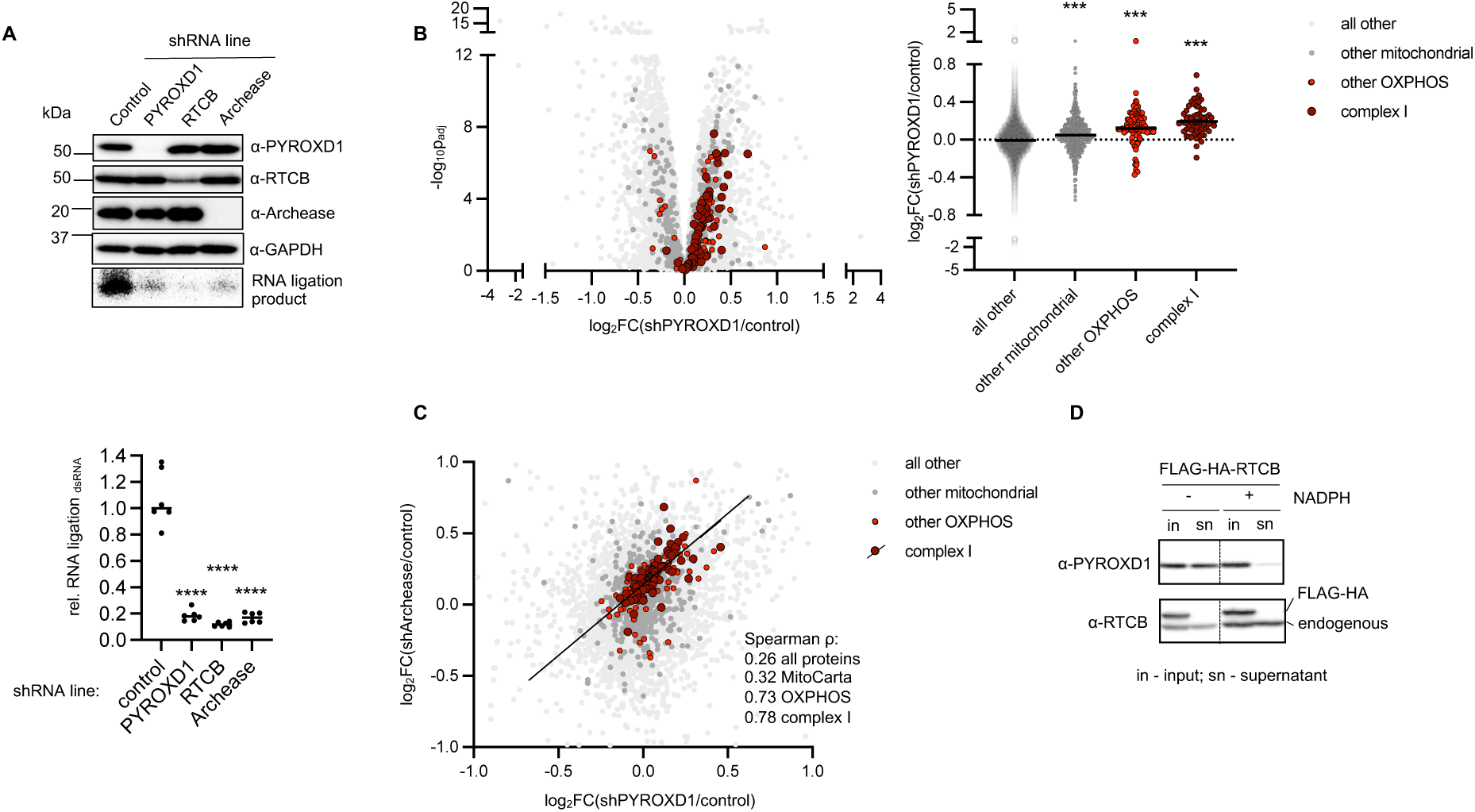
Cellular PYROXD1 is largely bound to the tRNA-LC, whose inactivation phenocopies loss of PYROXD1 in mitochondrial deregulation. **A.** PYROXD1, RTCB and Archease were silenced for 7 days in HeLa cells using genetically encoded, doxycycline-induced shRNA, and dsRNA oligo ligation activity was measured in extracts and detected by urea PAGE and autoradiography. Protein levels were assessed by western blotting and the ligation efficiency was quantified relative to the control line. The experiment was conducted 6 times. **** p<0.0001 (student t-test). **B.** PYROXD1 was silenced as in (A) and the whole cell proteome changes were analysed using mass spectrometry. The experiment was conducted 5 times. Mitochondrial proteins were identified based on MitoCarta 3.0 database. *** p<0.001 (student t-test) **C.** The experiment was conducted as in (B), with silencing of Archease instead of PYROXD1. Linear fit for complex I subunits is displayed. **D.** FLAG-HA-RTCB was immunoprecipitated from HEK293 Flp-In cells upon 1 day of doxycycline-induced overexpression with NADPH spiked in the lysis buffer where indicated. RTCB and PYROXD1 were detected in the supernatant by western blotting.

We next analysed the proteomic alterations in HeLa cells depleted of PYROXD1. As recently reported in myoblasts where PYROXD1 had been eliminated by CRISPR-mediated knock-out^37^, proteins involved in oxidative phosphorylation (OXPHOS) were consistently upregulated (**Fig. 1B**, left). Complex I displayed a particularly strong effect, and non-OXPHOS (oxidative phosphorylation) mitochondrial proteins only displayed a minor change (**Fig. 1B**, right). We contrasted this observation with an analysis of mouse hearts upon tamoxifen-inducible PYROXD1 knock-out^19^, where we also observed a mitochondrial deregulation, albeit in the opposite direction (**Fig. S1D**). This is in line with the previously published discrepancy in proteomic footprint between cultured myoblasts and skeletal muscle^37^, highlighting the possible adaptive differences between cultured cells and the living tissues. Therefore, while the direction of change appears condition-dependent, these data establish deregulation of the respiratory chain as a recurrent footprint of PYROXD1 depletion across cell lines, tissues and species.

We next tested whether silencing of Archease, which like PYROXD1 is critical for RTCB activity^18,20^, causes similar deregulation. In contrast to non-specific proteomic changes, depletion of Archease and PYROXD1 showed a marked correlation in deregulation of OXPHOS (**Fig. 1C**). Although the mechanism remains mysterious, this observation directly links changes in OXPHOS with failed activation or protection of the tRNA-LC, and implies that the critical cellular phenotypes of PYROXD1 occur due to incomplete protection of RTCB.

We have previously revealed that the members of the tRNA-LC are by far the most prominent interactors of PYROXD1 (reproduced here in **Fig. S1E**)^19^, and that PYROXD1 and RTCB bind in an equimolar manner^21^. However, publicly available proteomic data^45^ suggest that cellular levels of PYROXD1 are typically order(s) of magnitude lower than RTCB (**Fig. S1F**), implying cellular excess of RTCB. To test if majority of PYROXD1 is bound to RTCB in the cells, we tested how much PYROXD1 is depleted from cell extracts upon immunoprecipitation of FLAG-HA-RTCB. Indeed, FLAG-HA-RTCB near-quantitatively depleted PYROXD1 despite close-to-endogenous overexpression levels (**Fig. 1D**). Such depletion is reliant on NADPH, the ubiquitous cellular cofactor supporting the interaction^19^. In contrast, most of RTCB remained in solution upon precipitation of FLAG-HA-PYROXD1, even in presence of NADPH (**Fig. S1G**). The imbalance is also reflected at the genetic level. In co-essentiality maps^46,47^, which identify genes with shared fitness effects across cell lines, tRNA-LC subunits rank highest for PYROXD1, while PYROXD1 ranks markedly lower for RTCB (**Fig. S1H**).

Together, these data imply that PYROXD1 near-exclusively associates with RTCB in cells, and that the key downstream cellular changes including the effects on mitochondria arise due to loss of RTCB protection, giving us impetus to test whether RNA ligation is affected by PYROXD1 variants found in patients.

### All patient PYROXD1 variants fail to fully sustain RNA ligation

Currently available systems are not suitable for screening of potentially subtle differences in the performance of PYROXD1 variants. Relatively long PYROXD1 half-life and the residual RNA ligation in cells depleted of PYROXD1 by shRNA technology (**Figs. 1A, S1A**) create experimental background that could mask any changes caused by the partially dysfunctional pathogenic proteins. Three systems have been used to assess PYROXD1 variants so far, each with limitations: (i) yeast complementation assay^30,31,34^, limited by the absence of RTCB in fungi^19^; (ii) analysis of UPR in patient fibroblasts^29^, constrained by the robustness of XBP1 splicing (**Fig. S1C**) and by limited variant combinations and patient materials; and (iii) *in vitro* reconstitution^19,21^, which lacks cellular context.

To overcome these limitations, we endogenously tagged PYROXD1 in HeLa cells with a degradation (FKBP12^F36V^, here: deg-) tag^48,49^, allowing for speedy and quantitative clearance of PYROXD1. While endogenous tagging somewhat lowered PYROXD1 levels, the addition of degradation (dTAG^V-1^, here: dTAG) drug caused a complete loss of PYROXD1 and corresponding abrogation of short dsRNA oligonucleotide ligation in extracts (**Fig. 2A**). Both endogenous reactions: ligation of pre-tRNA fragments (**Fig. 2B**) and *XBP1* mRNA splicing (**Fig. S2A**), were dramatically impaired, ultimately resulting in loss of cellular viability (**Fig. 2C**). Similar dynamics of viability loss and the corresponding drop in tRNA maturation were observed upon clearance of the “core” tRNA-LC subunit DDX1^16^ (**Fig. S2B**), known for its specific involvement in tRNA splicing^50^. This implies that cellular essentiality of PYROXD1 likely stems from its role in sustaining pre-tRNA fragment ligation by the tRNA-LC.

**Figure 2.**
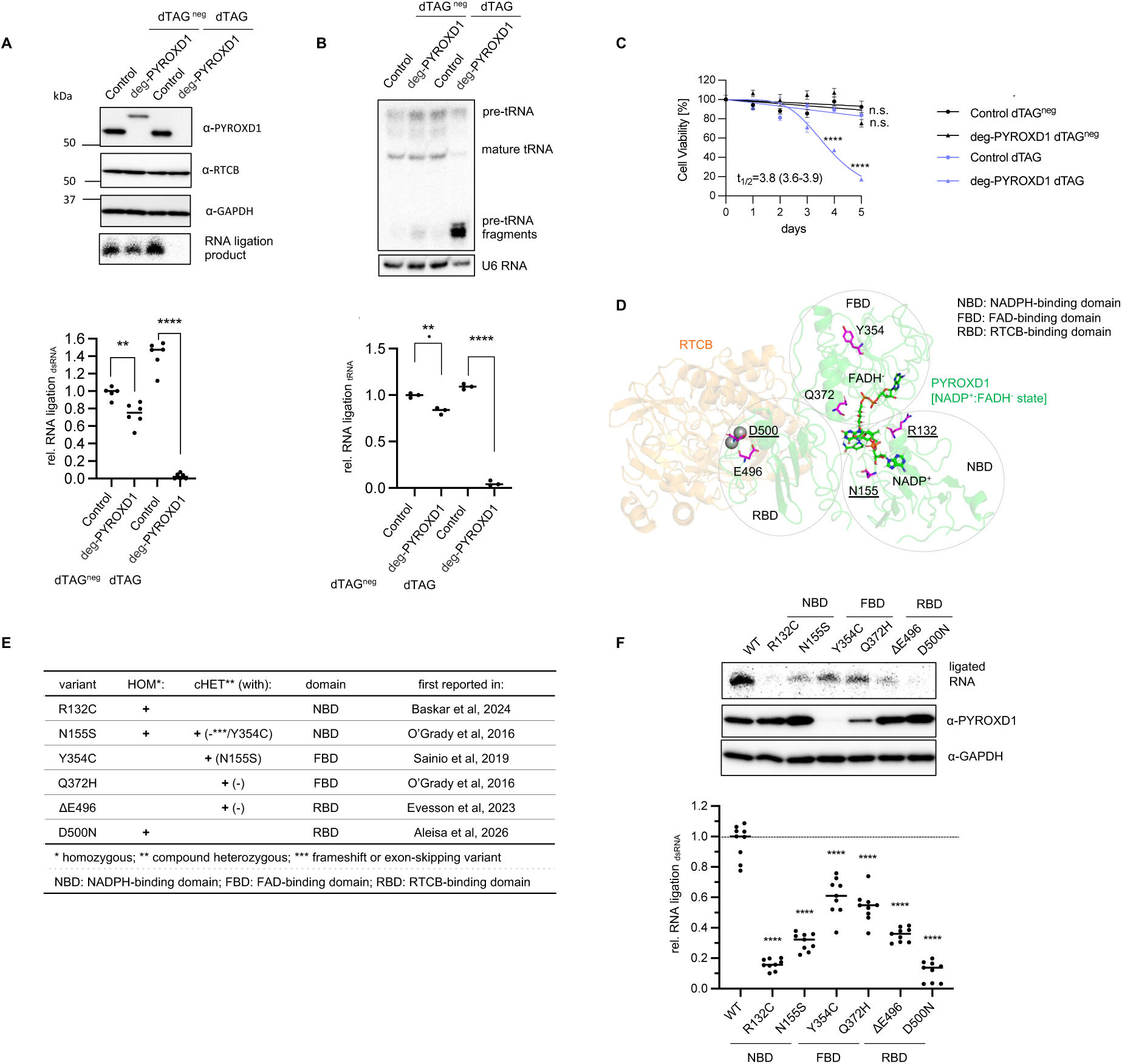
Coupling of degron-mediated silencing of PYROXD1 with constitutive expression of its patient variants allows to estimate their effects on RNA ligation. **A.** PYROXD1 was endogenously tagged with FKBP12^F36V^ (here: deg-) tag in HeLa cells, and depleted from cells for 3 days upon addition of degradation (dTAG^V-1^, here: dTAG) drug. Inactive degrader mimic (dTAG^V-1_neg^, here: dTAG^neg^) was used as a negative control. dsRNA oligo ligation activity was measured in extracts and detected by urea PAGE and autoradiography. Protein levels were assessed by western blotting and the ligation efficiency was quantified relative to control line treated with dTAG^neg^. The experiment was conducted 6 times. **** p<0.0001 (student t-test) **B.** Cells were treated as in (A) and pre-tRNA fragment ligation was assessed in the extracted RNA by northern blotting for Tyr^GTA^, and quantified relative to control line treated with dTAG^neg^. The experiment was conducted 3 times. ** p<0.01. **** p<0.0001 (student t-test) **C**. deg-PYROXD1 and control cells were treated for five days with dTAG or dTAG^neg^ and viability was monitored by AquaBluer staining. N=23. **** p<0.0001. Extra sum-of-squares F test was used for the whole curve comparison. **D**. Previously published cryo-EM structure of PYROXD1 in charge-transfer complex state in complex with RTCB (PDB: 8ORJ). Six known amino acids affected in reported single amino acid variants are highlighted in magenta, and those replaced or deleted in homozygous patients are underlined. Three domains of PYROXD1 (FBD: FAD-binding domain; NBD: NADPH-binding domain; RBD: RTCB-binding domain) are encircled. **E**. Overview of single amino acid patient PYROXD1 variants with publications first reporting them. + indicates at least one patient identified harbouring the designated variant in homozygous (HOM) or compound heterozygous (c-HET) genotype. **F**. FLAG-HA-PYROXD1 variants were stably integrated into deg-PYROXD1 line described in (A), resulting in constitutive expression. Endogenous PYROXD1 was depleted for 3 days with dTAG drug and dsRNA ligation was assessed in extracts followed by detection by urea PAGE and radiography, and quantified relative to WT. PYROXD1 levels were estimated by western blotting. The experiment was conducted 6 times. *** p<0.001 **** p<0.0001 (student t-test)

Both shRNA-mediated silencing (**Fig. S1A**) and degron-mediated clearance of PYROXD1 (**Fig. 2B**) impaired ligation of pre-tRNA fragments, evident in their cellular accumulation. Only quantitative, degron-mediated removal additionally caused a marked loss of mature tRNAs (**Fig. S2C**, top right) and impaired viability (**Fig. 2C**). The residual PYROXD1 in shRNA cells (**Fig. S2C**, bottom left) is therefore sufficient to sustain their mature tRNA levels and viability. This illustrates the robustness of mechanisms sustaining cellular mature tRNA pools, and emphasises the need for quantitative removal of endogenous PYROXD1 when assessing the full scope of its cellular phenotypes.

With a rapid and background-free system for depleting endogenous PYROXD1 at hand, we next focused on dissecting the effects of PYROXD1 variants found in patients. So far, six single amino acid variants of PYROXD1 were identified in individuals with PYROXD1 myopathy^29–31,34,36^. The variants affect largely conserved residues (**Fig. S2D**) that are scattered across the three domains of PYROXD1 (**Fig. 2D**): the FAD-binding domain (FBD, with the flavin-based prosthetic group), the NADPH-binding domain (NBD, responsible for accommodating the cofactor NADPH), and the RTCB-binding domain (RBD, providing the platform for interaction with RTCB primarily through the C-terminal tail^21^; previously called “C-terminal domain”^19^). Three variants were so far identified in patients at homozygosity^30,31,36^, while the others are found in compound heterozygous forms^29,34^, often with presumably deleterious frameshift or exon skipping variants (**Fig. 2E**).

We generated cell lines constitutively expressing WT PYROXD1 or the variants of interest whose open reading frames are IRES-coupled to GFP^51^ to ensure homogenous expression rates (**Fig. S2E**). We then removed endogenous deg-PYROXD1, and assessed RNA ligation in extracts. All variants failed to fully sustain RNA ligation (**Fig. 2F**).

We next assessed RNA ligation with endogenous substrates. *XBP1* mRNA splicing appeared robust across variants (**Fig. S2F**), in line with the observation that only full removal of PYROXD1 has a major impact on UPR. In contrast, endogenous pre-tRNA fragment ligation was affected in all lines (**Fig. S2G**), with two variants displaying the lowest RNA ligation in extracts also accumulating the most pre-tRNA fragments. Moderate and mild variants could not be finely differentiated by northern blotting, likely due to the long half-life of tRNA species in cells. To overcome this, we performed pre-tRNA splicing assays in extracts using radiolabelled pre-tRNAs, and obtained a pattern indistinguishable from the ligation assays with dsRNA (**Fig. S2H**, compare with **Fig. 2F**). We therefore conclude that for engineered variant cell lines with inducible depletion of endogenous PYROXD1, RNA ligation in extracts provides the most sensitive readout for enzymatic RTCB activity and thus PYROXD1 efficacy as its protector.

This analysis of RNA ligation in the extracts of cells expressing individual variants appeared to align with clinical data. For example, all three variants found in homozygous patients ranked at the bottom of RNA ligation sustenance, with the most severe variant D500N^31^ displaying the lowest RTCB support. Patient PYROXD1 variants therefore converge on impaired RNA ligation in a manner apparently consistent with pathogenicity.

### Variants in FAD-binding domain cause PYROXD1 instability

All PYROXD1 variants found in myopathy patients fail to support RNA ligation (**Fig. 2F**, top bar). Two of them, Y354C and Q372H (both within the FBD) are expressed at lower levels in our degron system (**Figs. 2F** middle bar**, S2E,H**), suggesting that their defect may at least in part arise from loss of protein stability rather than from a specific disruption of the PYROXD1–RTCB interplay.

To test if inherent faults in protein folding trigger lower expression, we employed thermal shift assays. We first determined melting point of WT PYROXD1 (**Fig. S3A**), followed by the analysis of the patient variants (**Fig. 3A**). Regardless of the presence of cofactor NADPH, the least stable variant was Q372H, supporting its decreased expression levels. Q372 resides in the 358-376 loop within FAD binding domain, with residues both upstream and downstream participating in FAD binding (**Fig. S3B**). Failure of any tested replacement of Q372 to provide WT protein stability (**Fig. S3C**) implies the critical role of Q372 in stabilising the 358-376 loop at the core of FBD, and potentially in binding of the prosthetic group.

**Figure 3.**
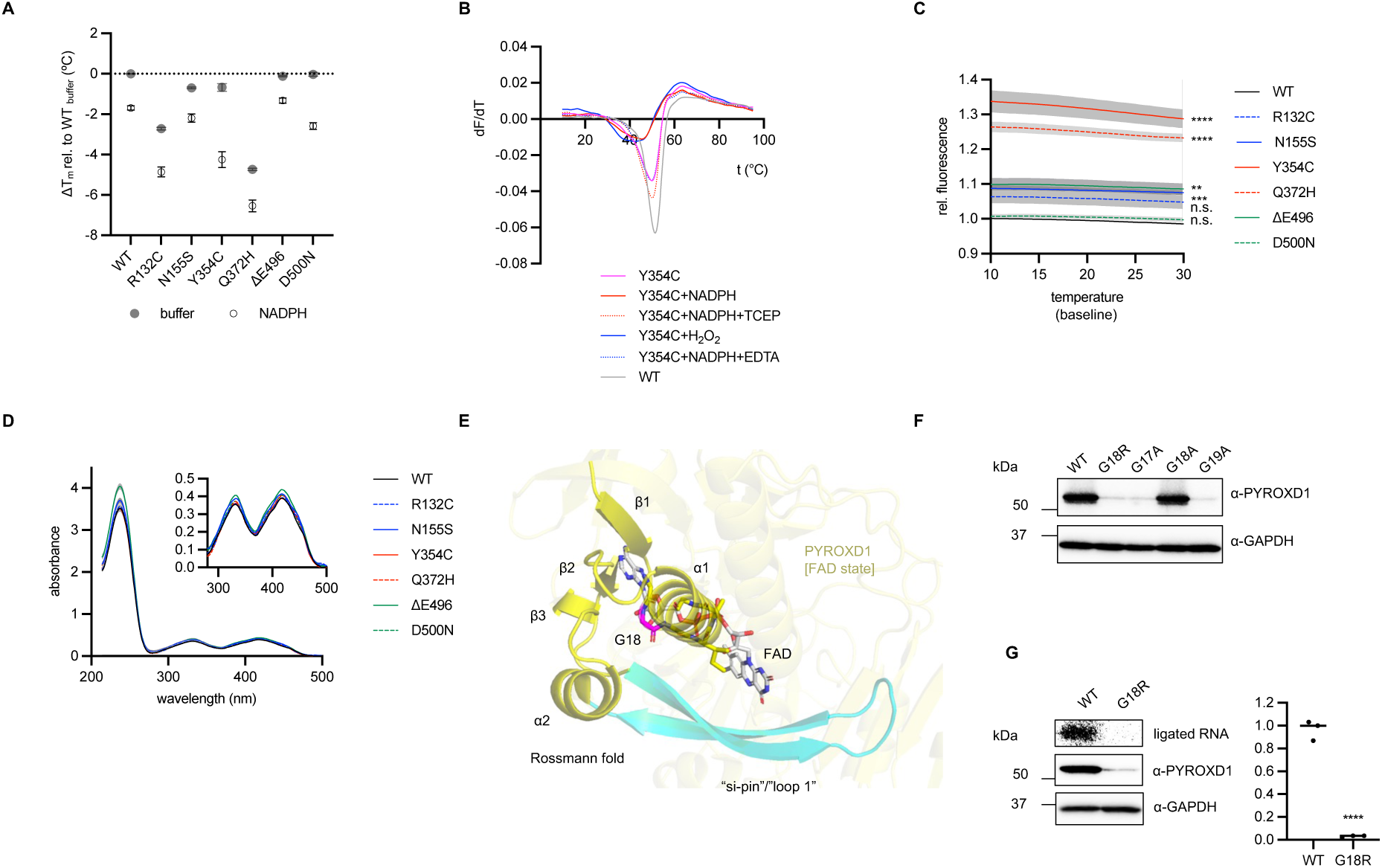
FAD binding domain variants, including a potentially novel pathogenic replacement G18R, display lower protein stability. **A.** Recombinant PYROXD1 variants were expressed in *E. coli* from pGex-6p-1 plasmid and purified from using GST-affinity capturing, tag cleavage and size exclusion. The proteins were incubated with 1 mM NADPH or buffer for 1 h at room temperature as designated. Melting points were assessed by SYPRO Orange thermal shift assay as centroids of melting curve derivatives. N≥9. Each measurement is conducted with a separate aliquot from at least two independent preparations. Statistical significance (student t-test). Buffer (relative to WT) R132C **** N155S **** Y354C ** Q372H **** ΔE496 ** D500N n.s., NADPH (relative to NADPH-treated WT): R132C **** N155S n.s. Y354C **** Q372H **** ΔE496 n.s. D500N ** **B**. Y354C PYROXD1 variant, expressed and purified as in (A), was incubated for 1 h at room temperature as designated (NADPH: 1 mM; H2O2: 250 μM; EDTA: 0.5 mM), and the melting was assessed by SYPRO Orange assay. Derivative curves are displayed. **C**. Samples were treated as in (A). Pre-unfolding baseline range of non-derivatised thermal shift curves is displayed. N≥9, average value and SEM are displayed. Extra sum-of-squares F test was used for the whole curve comparison. ** p<0.01 *** p<0.001 **** p<0.0001 **D**. Ultraviolet-visible spectra are recorded from the recombinant preparations of patient PYROXD1 variants. N≥6. Each measurement is conducted with a separate aliquot from at least two independent preparations. Average value and SEM are displayed. **E**. Previously published PYROXD1 structure in FAD form, PDB: 6ZK7. PYROXD1 contains an atypical monomeric Rossmann fold interrupted by a pin unique among flavoproteins. **F**. HeLa cells, with PYROXD1 endogenously tagged by FKBP12^F36V^ tag were depleted of endogenous PYROXD1 for 3 days after being transiently transfected with Lipofectamine 3000 with plasmid expressing FLAG-HA-PYROXD1 constructs of interest. Protein levels were assessed by western blotting. Experiment is conducted three times and representative image is displayed. **G**. PYROXD1 was endogenously tagged with FKBP12^F36V^ tag in HeLa cells, and FLAG-HA-G18R or WT PYROXD1 were stably integrated, resulting in constitutive expression. Endogenous PYROXD1 was depleted for 3 days with dTAG drug and dsRNA ligation was assessed in extracts followed by detection by urea PAGE and autoradiography, and quantified relative to WT. PYROXD1 levels were estimated by western blotting. Experiment was conducted 3 times. **** p<0.0001 (student t-test).

The least expressed Y354C variant (**Fig. 2F, S2E** bottom, **S2H**) showed only a minor melting point defect, but displayed the steepest melting point drop upon incubating with NADPH (**Fig. 3A**). A closer inspection of the melting curve derivative revealed a low-stability species already partially melted at body temperature. This species could however be rescued with TCEP and EDTA, and was also generated when H₂O₂ was used instead of NADPH (Fig. 3B). This implies that NADPH-triggered destabilisation is in fact caused by H₂O₂, which accumulates due to slow NADPH oxidation catalysed by PYROXD1^19^, and not by nucleotide binding itself. The effect is particularly pronounced for Y354C. Low cellular expression levels of Y354S and Y354D, which could mimic oxidised cysteine states, is consistent with potential oxidation of C354 as a cause of Y354C destabilisation (**Fig. S3D**).

However, even in the presence of NADPH, Y354C does not display the lowest melting point among variants (**Fig. 3A**) as expected based on cellular expression. This implies that additional mechanisms may contribute to cellular clearance of Y354C.

Exploring other thermal shift parameters, we observed that the fluorescence baseline markedly increased in Q372H and in particular in Y354C variant (**Fig. 3C**), a pattern preserved in presence of NADPH (**Fig. S3E**). The major effect found in Q372H and Y354C could not be explained by varying protein concentration, as evidenced by largely overlapping UV-visible spectra across variant preparations (**Fig. 3D**) and comparable amounts on SDS-PAGE (**Fig. S3F**). We therefore propose that variants in FBD alter PYROXD1 conformation, potentially exposing hydrophobic patches that make the protein prone to cellular removal. Such conformational change may occur due to altered FAD binding, supported by the position of Y354 at the core of FBD, interplaying with its central 376-394 helix (**Fig. S3G**). The first helix residue W376 is critical for stacking with FAD in NADPH-free state^19,21^, and also an endpoint of Q372-containing 358-376 loop (**Fig. S3B**). Y354C and Q372H thus have altered conformation under baseline conditions, undergoing additional destabilisation due to cysteine oxidation particularly prominent in Y354C.

The role of FAD for PYROXD1 stability gained further prominence when we focused on a potentially novel G18R variant, currently only published in the context of a conference abstract^38^. G18 is a part of the non-canonical Rossmann fold^52^ of PYROXD1, unique for its lack of dimerisation and the “si-pin” (called loop 1 within PYROXD1 in NADP^+^:FADH^-^form^21^) inserted between the second α-helix and the third β-strand^19^ (**Fig. 3E**). <u>G18</u> specifically resides within G<u>X</u>GXXG Rossmann fold motif^53^ (G<u>G</u>GIAG in human PYROXD1) widely utilised by enzymes to bind FAD and other dinucleotides, where it surrounds the pyrophosphate group between the two nucleotides^54^ (**Fig. S3H**). In line with the G<u>X</u>GXXG consensus, mutational analysis of the residues 17-19 matches the invariability of Gs at positions 17 and 19, and tolerance for mild replacement at position 18 (**Fig. 3F**). However, the introduction of a bulky, charged residue R18 was not permitted, conveying the variant its likely pathogenicity.

Attempts at purifying G18R constructs resulted in extremely low yields (**Fig. S3I**) insufficient for reliable characterisation *in vitro*. This is in line with our previous finding that the WT PYROXD1 is fully loaded with FAD^19^ and our failures to purify artificial constructs with replacements in FBD^21^, as well as with the observation that despite markedly different yields, only the fully FAD-loaded fraction of each variant could be successfully purified (**Fig. S3J**).

The drastic effect of G18R replacement was also observed in a cell line constitutively expressing the variant on top of inducible depletion of the endogenous protein. Here, the G18R line displayed diminished RNA ligation (**Fig. 3G**) coupled with loss of mature tRNAs (**Fig. S3K**) and *XBP1* mRNA splicing (**Fig. S3L**), so far only observed in cells fully lacking PYROXD1 (**Fig. 2B**). We therefore assume that the remaining PYROXD1 traces surviving massive destabilisation are improperly folded. G18R would hence likely phenotypically mimic frameshift or exon skipping PYROXD1 variants, triggering complete loss of functional protein, and should only be found in heterozygous individuals.

Together, these data revealed that the two known patient variants in FBD, as well as the potentially novel G18 replacement, display lower expression levels, likely consequent to the structural changes caused by impaired FAD binding.

### Defective redox switch in NADPH-binding domain variants hinders RTCB association

Controlled oxidation of NADPH is central to the protective role of PYROXD1 towards RTCB^19^ (**Fig. 4A**). Rather than displaying a *bona fide* NADPH oxidase activity, PYROXD1 reacts with NADPH to generate a NADP^+^:FADH^-^charge transfer complex (CTC) (step 1, “on” reaction), triggering unstructuring of loop 1 and allowing RTCB binding^21^. CTC ultimately gets oxidised by O_2_ (step 2, “off” reaction), allowing Archease to bind to RTCB upon PYROXD1 dissociation. The CTC-bound state is thus the functionally critical form of PYROXD1 that binds and protects RTCB.

**Figure 4.**
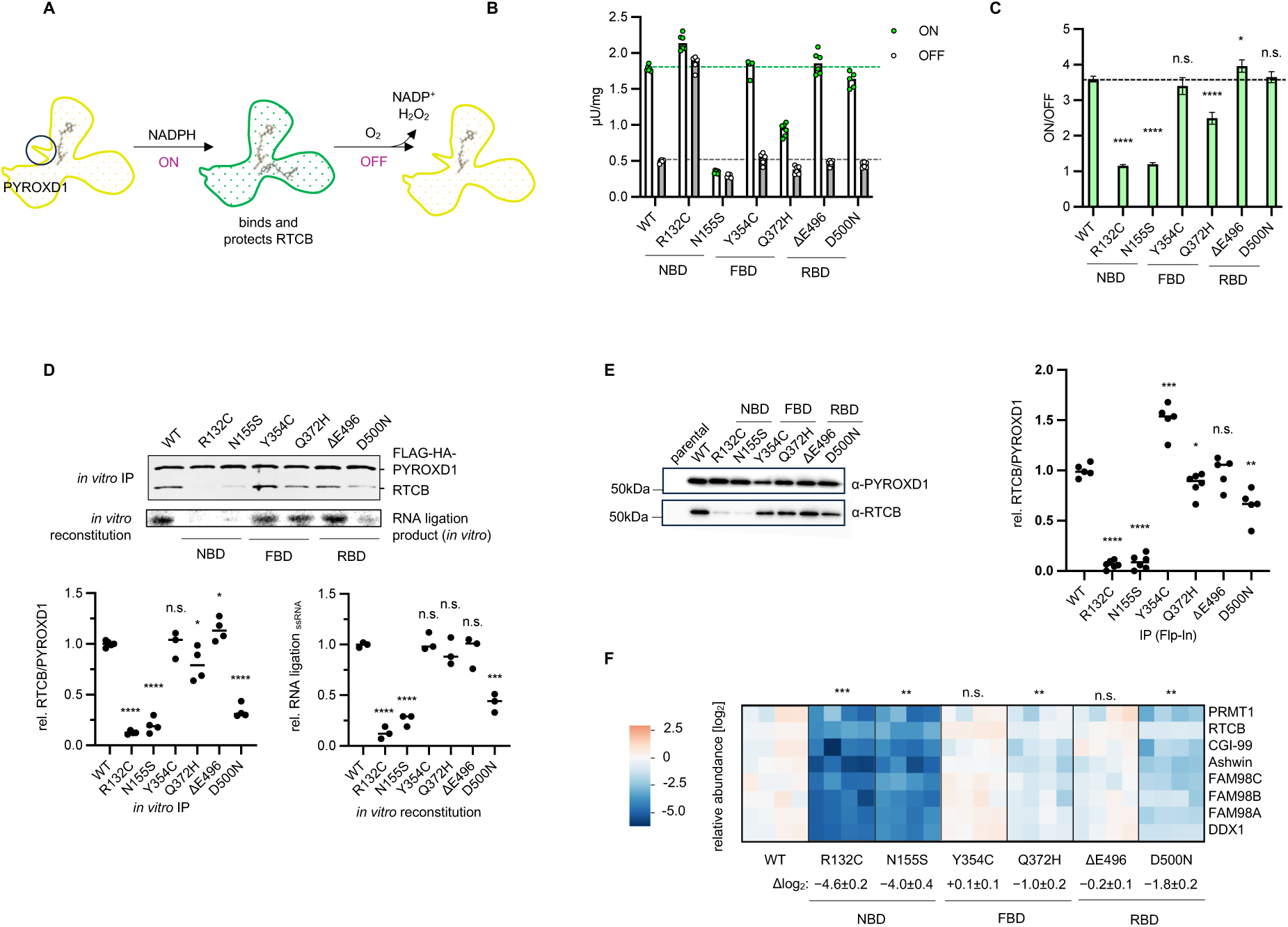
Variants in NADPH-binding domain display defective flavin-switch and consequently lowered binding of RTCB. **A.** Schematic representation of the controlled oxidation of NADPH by PYROXD1. Only upon formation of the NADP^+^:FADH^-^ charge transfer complex (CTC) loop 1 becomes disordered, removing steric hindrance (encircled left) and allowing binding and protection of RTCB. **B**. Recombinant PYROXD1 variants were expressed in *E. coli* from pGex-6p-1 plasmid and purified from using GST-affinity capturing, tag cleavage and size exclusion. Proteins were incubated with NADPH together with either artificial quinoid substrate menadione or buffer (where O_2_ serves as electron acceptor) to assess the on and off reaction rates during PYROXD1 reaction cycle defined in (A). Reaction was monitored for 45 minutes and NADPH oxidase (buffer) or dehydrogenase (menadione) activity was determined from the linear range of the absorbance loss at 340 nm. Activity unit is defined as the amount of enzyme that converts 1 μmol of NADPH per minute and the protein concentration was determined with Bio-Rad Protein Assay. N=6. Statistical significance: ON R132C **** N155S **** Y354C n.s. Q372H **** ΔE496 n.s. D500N *, OFF R132C _**** N155S **** Y354C n.s. Q372H ** ΔE496 n.s. D500N * (student t-test) **C**. Estimated ON/OFF ratios were determined from the data displayed in (B) and the experimental error and statistical significance were determined by bootstrap resampling. **** p<0.0001 * p<0.05 **D**. Recombinant FLAG-HA-PYROXD1 recombinant proteins were expressed in *E. coli* from pGex-6p-1 plasmid and purified from using GST-affinity capturing, tag cleavage and size exclusion. Proteins were incubated with the recombinant tRNA-LC (4 subunits, untruncated) in presence of NADPH, immunoprecipitated and washed with NADPH-containing buffer (top). FLAG-HA-PYROXD1 and RTCB were visualised by silver staining, and RTCB/PYROXD1 ratio was quantified and displayed relative to WT (bottom left). **** p<0.0001 * p<0.05 (student t-test). Recombinant tRNA-LC (4 subunits, untruncated) was incubated with NADPH, H2O2, Archease and PYROXD1 for 1 h at room temperature. Afterwards, it was mixed with TCEP, ATP, GTP and radiolabelled ssRNA oligo. RNA circularisation was visualised by autoradiography (middle) upon separation with urea PAGE, and quantified relative to WT (bottom right). **** p<0.0001 *** p<0.001 (student t-test). **E**. FLAG-HA-PYROXD1 was immunoprecipitated from HEK293 Flp-In cells upon 1 day of doxycycline-induced over-expression, with lysis buffer and wash buffers supplemented with NADPH. Immunoprecipitated PYROXD1 and RTCB were visualised by western blotting (left) and RTCB/PYROXD1 ratio was quantified and displayed relative to WT (right). Experiment was conducted five times. **** p<0.0001 *** p<0.001 ** p<0.01 * p<0.05 (student t-test). **F**. FLAG-HA-PYROXD1 variants were immunoprecipitated as in (E). The immunoprecipitates were analysed by mass spectrometry. PYROXD1 and the tRNA-LC components, together with the associated modifier PRMT1, are displayed. Experiment was conducted 4 times. Δlog2 annotates average deregulation of the designated subunits, and error represents SEM with N=4. *** p<0.001. ** p<0.01 (student t-test)._

Since the “off” reaction is rate-limiting with O_2_ as the electron acceptor^19^, the “off” rate approximately reflects the overall rate of NADPH consumption. To be able to also measure the “on” rate, we performed the same reactions with artificial quinoid substrate menadione, which instantaneously oxidises the CTC and makes step 1 rate-limiting^19^. Hence, by measuring NADPH turnover in presence of menadione and O_2_, we could gauge the relative “on” and “off” rates of CTC generation and spending, respectively (**Fig. 4B**).

Two variants, N155S and R132C, displayed striking changes in NADPH oxidation kinetics. N155S shows a slower “on” rate, in line with previous pre-steady-state measurements and the role of N155 in positioning NADP(H)^19^ (**Fig. S4A**). To test whether this is due to loss of the hydrogen bond between the side-chain amide of N155 and NADPH, we transiently expressed N155G and N155A, which abolish the side-chain contacts. Both constructs however successfully support RNA ligation (**Fig. S4B**), implying that the altered N155S interaction is more detrimental than complete loss of the interaction. Consistently, N155G mice display no myopathy phenotype and no mitochondrial deregulation^37^, further strengthening the link between RNA ligation and the key phenotypic consequences. N155G interestingly displays an elevated “on” rate (**Fig. S4C**, top) and “on”/”off” ratio (**Fig. S4C**, bottom), implying that increasing CTC half-life beyond WT levels does not create a super-protector that markedly boosts RTCB support and leaves a distinct proteomic footprint on the cells.

In contrast to N155S and N155G, R132C displayed increased “off” rate, i.e. increased reactivity in presence of oxygen. This makes R132C the first “hyperactive” variant found in a myopathy patient with an increased turnover, and could be attributed to the role of R132 in the interaction network that restricts access of O_2_ to the CTC by positioning “loop 2” above FAD (**Fig. S4D**). PYROXD1 thus needs to operate within a defined NADPH oxidase activity range and deviation in any direction may cause pathogenicity, as previously speculated based on an artificial construct^19^.

While “too slow” step 1 and “too fast” step 2 in N155S and R132C respectively appear to have contradictory effects on the overall NADPH turnover, they likely converge in lowering the time PYROXD1 spends in CTC-containing, RTCB-binding form. We therefore propose that, rather than individual reaction rates, the lowered on/off ratio (**Fig. 4C**) should be considered a more universal predictor of PYROXD1 dysfunctionality in terms of RTCB protection, reflecting inability of PYROXD1 to undergo controlled conformational changes.

In addition to R132C and N155S, the previously discussed Q372H variant also displays lowering of the “on/off” ratio, albeit to a milder extent. The loop 358-376 may help in positioning of NADP(H) despite it largely residing within FBD (**Fig. S4E**). Hence, defective NADPH switch could play a role in Q372H pathogenicity alongside its lowered stability in cells.

If “on”/”off” ratio reflects the time PYROXD1 spends in CTC form, a drop in “on/off” value should result in lowered binding to RTCB. To test this, we performed *in vitro* IP using purified recombinant proteins and NADPH. In line with NADPH switch analysis, both R132C and N155S displayed a stark drop in RTCB binding, with Q372H displaying a milder effect (**Fig. 4D**, top, quantified bottom left). The only exception not predicted by the altered “on”/”off” ratio was D500N (see below). Loss of RTCB binding across variants resulted in proportionally lowered protection of the tRNA-LC in the previously established *in vitro* reconstituted system^19^, where the tRNA-LC and Archease were co-incubated with PYROXD1, NADPH and H_2_O_2_ before adding substrates for RNA ligation (**Fig. 4D**, middle, quantified bottom right).

This pattern was also largely reflected in the immunoprecipitates from Flp-In HEK293 lines over-expressing FLAG-HA-tagged PYROXD1 variants, confirming that the inherent features of PYROXD1-RTCB interplay are retained in the context of cell extracts (**Fig. 4E**, **S4F**). Y354C appeared to be an exception, with an apparently elevated RTCB/PYROXD1 ratio in western blot quantification (**Fig. 4E**, right). However, normalisation of co-immunoprecipitated RTCB to the FLAG-PYROXD1 bait is only reliable when bait levels are comparable across samples, which is not the case for the poorly expressed Y354C (see FLAG-HA-PYROXD1 variant levels in **Fig. 4E**). To overcome the limited dynamic range of western blot quantification, we analysed the variant immunoprecipitates by mass spectrometry (**Figs. 4F**, **S4G**), revealing that the entire tRNA-LC, together with the previously identified interactor PRMT1^55^, follows the same pattern as RTCB, in line with RTCB being the anchor for PYROXD1 binding and implying that PYROXD1 variants similarly affect all cellular forms of the tRNA-LC.

### Two variants target the PYROXD1 C-tail, a unique sequence evolved for RTCB binding

Two variants remained mechanistically mysterious after the analysis of expression levels and NADPH-mediated switch: ΔE496 and D500N. Both variants are located within the RBD of PYROXD1, specifically in the C-tail, a sequence only found in PYROXD1 and serving as a primary platform for binding of RTCB^21^ (**Figs. 5A, S5A**).

**Figure 5.**
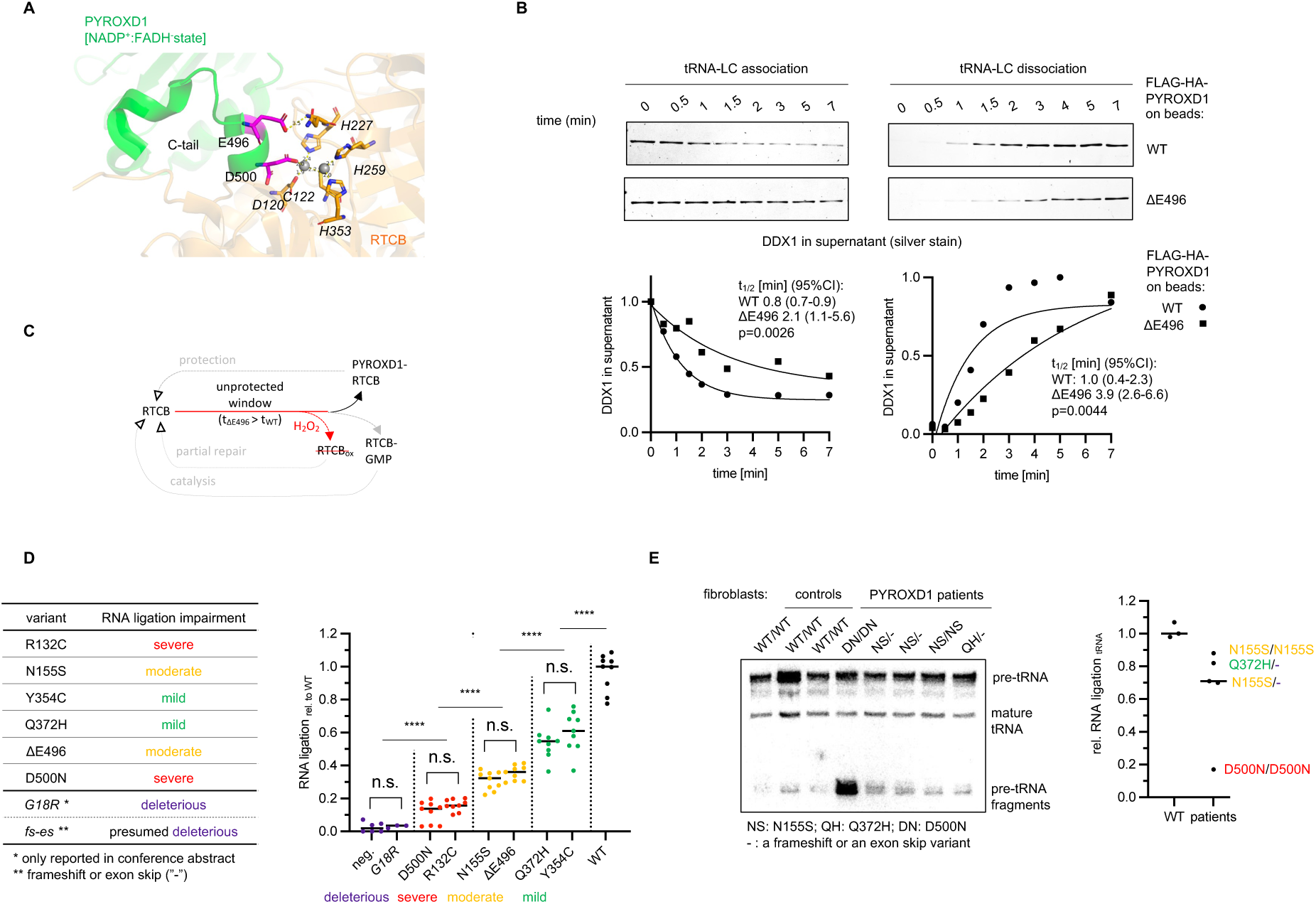
RTCB-binding domain variants have most specific impairments and inspire variant classification with predictive power. **A**. Both RTCB-binding domain variants affect C-tail, unique stretch of amino acids (UniProt peptide search; BLASTP score PYROXD1 E=4·10^-5^ vs best non-PYROXD1 hit E=1.1·10^-1^) inserted into RTCB active site, in previously published structure (PDB 8ORJ). D500 residue coordinates metal ions also bound by the active site cysteine residue in RTCB. **B**. Recombinant FLAG-HA-PYROXD1 were expressed in *E. coli* from pGex-6p-1 plasmid and purified from using GST-affinity capturing, tag cleavage and size exclusion. The protein was bound to the anti-FLAG beads in the presence of NADPH, and the recombinant tRNA-LC (4 subunits, untruncated) was subsequently added. The loss of the tRNA-LC due to association with immunoprecipitated PYROXD1 was monitored by periodic aliquoting from the supernatant (top left); The recombinant FLAG-HA-PYROXD1 and the tRNA-LC were co-incubated with anti-FLAG beads in the presence of NADPH, washed once with NADPH, and the buffer was exchanged for NADPH-free solution. The increase in tRNA-LC amount in the supernatant was monitored by periodic aliquoting (top right). In both panels, proteins were visualised by silver staining. DDX1 was represented and used for quantification (bottom) due to best staining among the tRNA-LC subunits because of its large size. Extra sum-of-squares F test from nonlinear regression was used to compute statistical differences. **C**. Upon every PYROXD1 dissociation, completion of catalytic cycle or repair by thioredoxin, RTCB enters a period of vulnerability to oxidation, whose length is determined by the speed of association of PYROXD1. **D**. Patient PYROXD1 variants were classified to severe, moderate and mild depending on the extent of their failure to support RNA ligation (based on Fig. 2F). Variants within each group displayed no statistical difference, while the three groups robustly clustered separately from each other (right). **** p<0.0001. Combination of categories present in the two alleles is a predictor of myopathy onset (see Fig. S5H). **E**. RNA was extracted from the primary fibroblasts derived from healthy individuals and PYROXD1 patients, and pre-tRNA fragment ligation was assessed through northern blotting against TyrGTA. Each dot represents a line derived from a different individual. Colour coding is used as in (D). Statistical analysis: WT vs all variants * (p<0.05); WT vs variants with at least one moderate or mild allele ** (p<0.01) (student t-test assuming unequal variances). The severe homozygote D500N inflates the variance of the patient group and may segregate to a distinct severe homozygous group in the future.

ΔE496 is particularly puzzling as it reveals no apparent major defect in any *in vitro* analyses so far, despite providing lowered protection of RTCB in cells (**Figs. 2F, S2H**). A control experiment provided the first hint towards its molecular defects: FLAG-HA-PYROXD1 immunoprecipitates from Flp-In lines in the absence of NADPH occasionally revealed trace amounts of RTCB, which was only consistently detected in the FLAG-HA-ΔE496 line (exemplified in **Fig. S5B**). In contrast, no RTCB was observed in NADPH-free *in vitro* IPs with recombinant proteins (**Fig. S5C**, left). However, when we included NADPH in the binding buffer to mimic the cofactor present in the cell extracts but left it out during the washes, immunoprecipitates contained some RTCB, in particular when FLAG-HA-ΔE496 PYROXD1 was used as a bait (**Fig. S5C**). This suggested that loss of E496 may alter binding kinetics of PYROXD1 to RTCB. This was directly confirmed when we tested association (**Fig. 5B**, left) and dissociation (**Fig. 5B**, right) of the tRNA-LC to and from previously immunoprecipitated FLAG-HA-PYROXD1, respectively. Both processes were markedly delayed in ΔE496 variant despite comparable tRNA-LC amounts in the input and in the final immunoprecipitate (**Fig. S5D**).

ΔE496 therefore slows both association and dissociation. Since the C-tail occupies the same site on RTCB as Archease^20^ and nucleic acid substrate^56^, delayed dissociation could in principle inhibit RTCB by clogging the active site. However, neither an excess of recombinant ΔE496 *in vitro* (**Fig. S5E**), nor over-expressed FLAG-HA-ΔE496 in control cells (**Fig. S5F**) inhibited ligation. This suggests that delayed dissociation does not turn PYROXD1 into a competitive inhibitor of RTCB.

With inhibition excluded, we next focused on the other kinetic defect: delayed association. We propose that delayed PYROXD1 binding prolongs the unprotected window that opens each time free RTCB appears (**Fig. 5C**): upon PYROXD1 dissociation, after repair of oxidised RTCB^57^, and after conclusion of a catalytic cycle. These three routes simultaneously operate only in cells, and their accumulated effects are thus not easily recapitulated by *in vitro* assays. ΔE496 thus defines a distinct pathogenicity mechanism where RTCB protection is impaired by kinetic defects rather than by loss of activity or binding interface.

In contrast, the D500N variant displayed lowered binding to RTCB upon immunoprecipitation (**Figs. 4D-F, S4F-G**). The D500 residue directly coordinates catalytic metal ions in the RTCB active site (**Figs. 5A, S5A**), which we reported as critical for PYROXD1 binding^21^. Transient overexpression of D500A and ΔD500 on top of depletion of endogenous PYROXD1 caused similarly deleterious effects on RNA ligation (**Fig. S5G**), confirming that any change that weakens docking of C-tail to RTCB active site renders PYROXD1 largely incapable of sustaining RNA ligation.

Two variants, ΔE496 and D500N, affecting a sequence only known for binding of RTCB, and with no major negative effect on stability or redox-activity of PYROXD1, therefore selectively impede the interplay with the tRNA-LC.

### Specificity and severity of RNA ligation impairment align with clinical presentation

C-terminal D500N replacement in particular affects PYROXD1-RTCB interplay with surgical specificity at the core of the protective mechanism. The same variant displays the strongest clinical presentation, with the homozygous patient displaying congenital muscle weakness necessitating permanent life support^31^. This prompted us to test whether the relationship between RNA ligation defect and clinical presentation could be generalised.

We first classified the patient variants according to loss of RNA ligation activity (**Fig. 5D**). This includes the severe effect of a potentially new pathogenic variant G18R^38^, which is the first missense variant to be as deleterious for RNA ligation as total lack of PYROXD1 or presumably, frameshift or exon skipping variants. We then compared our ligation data with the reported disease onset in known patients. Patients with two severe or one severe/deleterious and one moderate variant displayed early onset; patients with two moderate or one mild and one severe/deleterious variant displayed mixed onset, while the only known person with one mild and one moderate variant had disease onset later in life (**Fig. S5H**).

The analysis so far has been based on the degron model, with each cell line only expressing a single variant of interest. Mechanisms of all disease variants however converge on lower occupancy of RTCB bound to PYROXD1. This implies that in heterozygous cells, each variant’s failures may contribute to loss of RNA ligation independently, in a potentially additive way. To test whether allelic combinations match outcomes based on the analysis of individual variants, we tested the primary patient fibroblasts for endogenous pre-tRNA fragment ligation (**Fig. 5E**). Cells from WT individuals clustered separately from all patient cells; furthermore, the patient cells with at least one allele with a mild or moderate variant^30^ showed markedly higher RNA ligation than the cells from the patient homozygous for the severe variant D500N^31^. This analysis implies that RNA ligation defects in individual variants can be the basis for estimating the expected defect in patient cells, and ultimately for anticipating the disease onset in new patients.

Such predictive power of RNA ligation defects, together with convergence of all variants on RTCB failure and the high specificity of RTCB-binding domain replacements, together present the impaired RNA ligation at the core of PYROXD1 myopathy.

## Discussion

This report reveals that the variants of PYROXD1 that cause severe myopathies in humans share a common pathomechanism, converging on defects in the ligation of RNA molecules. By dissecting molecular mechanisms affecting PYROXD1 variants, we achieved the first comprehensive analysis of pathogenic, protein coding variants of a factor directly involved in RNA ligation. As a sub-stoichiometric and more dynamic, yet functionally critical subunit of the tRNA-LC, PYROXD1 emerged as an indispensable tool to decipher the biomedical implications of defective RNA ligation upon discovery of rare myopathies caused by its multiple single amino acid variants.

Why did variants of PYROXD1, and not the other subunits, specifically rise in prominence as disease-causing factors? The closest case within the tRNA-LC is DDX1, where a single patient with microcephaly harbouring a protein-coding variant was described^27^, although no causative link or mechanistic pathogenicity basis has been established. Additionally, variants in the enhancer region proximal to DDX1, but also other genes including MYCN, were associated with cranial malformations^28^. Conditional deletion of DDX1 and artificially induced mature tRNA fragmentation cause cranial defects in zebrafish^28^, yet these interventions are not subtle and it remains unclear to which extent non-lethal enhancer variants would cause comparable phenotypes. Limited evidence thus implicates DDX1 in neuronal diseases, yet in the absence of protein-coding pathogenic variants, research on DDX1 often relies on its knock out, which may trigger pleiotropic effects due to its essentiality. This is also underscored by our analysis of PYROXD1. Cells with complete depletion display phenotypes (e.g. loss of mature tRNAs and cell viability) absent from the cells with partial gene silencing or from those expressing patient variants. Knock-out models of tRNA-LC subunits may thus not represent phenotypic consequence realistically expected in healthy or diseased tissues.

Other reports linking diseases to RNA ligation often affect upstream proteins. Variants of the subunits of tRNA splicing endonuclease complex (TSEN) are well-known causes of pontocerebellar hypoplasia (PCH)^58–60^. Most of the described variants affect TSEN assembly, and the observed changes in tRNA maturation were often mild and not proportional to clinical severity^58,61^. PCH is also caused by variants of CLP1^22–25^, a protein that associates with TSEN^62^, but also plays important roles in mRNA 3’end formation^63^. It is thus not clear whether tRNA splicing defects cause TSEN-linked PCH. tRNA-LC was recently revealed as the prime target of the novel degradation pathway dependent on FBXO31, whose variants cause neurodevelopmental disorder^64^. Proposed disease mechanism however includes neomorphic substrate recognition rather than loss of normal tRNA-LC turnover. In contrast, tRNA-LC was recently observed to be preferentially sequestered in protein aggregates in neuronal disorders such as fragile X-associated tremor/ataxia syndrome (FXTAS), due to the exceptionally glycine-rich tail of FAM98B subunit^26^. Here, the authors demonstrate defective pre-tRNA splicing, although the extent and mechanism of its contribution to pathogenesis remain to be established.

PYROXD1 myopathies stand apart as diseases caused by (multiple) pathogenic protein coding variants^29–31,34,36^ at the heart of the tRNA-LC. Prior to this work, it was unclear whether PYROXD1 has prominent moonlighting cellular pools independent of the tRNA-LC and if alterations in the tRNA-LC could cause PYROXD1-associated phenotypes such as deregulation of mitochondrial respiratory chain. Our analysis however implies that a vast majority of cellular PYROXD1 resides within the tRNA-LC, and that mitochondrial consequences of Archease silencing, the other subunit critical for function of the tRNA-LC, correlate with those triggered by PYROXD1.

This report reveals a striking concordance between the loss of RNA ligation activity in our cellular assays and the phenotypic presentation in patients. Our classification of PYROXD1 variants, directly rooted in the biochemical interplay with the tRNA-LC, provides the first mechanistically supported framework for prediction and explanation of how PYROXD1 variants impact its cellular function. Because the known PYROXD1 single amino acid variants show no evidence of mechanistic interplay or dominant-negative behaviour, their impacts on RNA ligation may be assumed to combine additively, extending the predictive power of our analysis to the biallelic combinations found in patients. Our determined average RNA ligation score of any two variants observed in patient cells was always below 50% relative to wild-type PYROXD1 (**Fig. S5I**). This defines a tentative disease threshold of ∼50% residual RNA ligation activity. Consistently, all known individuals with a single variant allele, where the WT allele keeps the average allele RNA ligation at or above 50%, are asymptomatic carriers, and no homozygous patients with the fully deleterious variant were described. In turn, our analysis suggests that an individual could not be homozygous for the G18R variant due to deleterious impact on PYROXD1 function and RNA ligation, as homozygosity would place them well below the viability threshold. The link between RNA ligation efficacy and phenotypic outcome is further strengthened by the mechanistic distinction between the pathogenic N155S variant, which fails to sustain RNA ligation, and the artificial N155G construct, which fully supports the tRNA-LC and causes no muscle phenotype in mice^37^.

Concordance between molecular phenotype (loss of RNA ligation) and disease presentation is particularly relevant for the variants in RTCB-binding domain. While protein destabilisation of FAD-binding domain variants or altered NADPH turnover found in NADPH-binding domain replacements would likely affect any plausible PYROXD1 function, C-tail of PYROXD1 is unique in human proteome and likely specifically critical for RTCB binding. C-tail variants^29^, including the most potent D500N replacement^31^, thus specifically affect the tRNA-LC with minimal consequences to wider PYROXD1 biochemistry.

Our work thus provides the first assay for assessing and explaining pathogenicity of PYROXD1 variants in a human cell line, which can easily incorporate novel variants or artificial constructs of interest. This comes in addition to establishing the feasible experimental tests such as on/off activity ratios, IP time courses and melting point tests that predict distinct modes of PYROXD1 biochemistry, each being sufficient to cause at least a partial RNA ligation defect. The thorough biochemical analysis further allows to predict the pathogenicity of variants, as demonstrated here with the potentially novel variant G18R, and which affected PYROXD1 stability in line with other FBD-affecting replacements. In addition to predicting and characterising the G18R variant, our work clarifies the status of additional variants. For example, the R132C, Y354C and D500N variants were all classified as “variant of uncertain significance” (VUS) upon their identification (ClinVar). This is despite both R132C^36^ and D500N^31^ being found in early-onset homozygous patients with severe clinical presentations. Here, we provide functional evidence establishing pathogenicity with a major impact on RNA ligation.

PYROXD1 is unique among the subunits of the tRNA-LC due to its partially conditional role. Protein coding variants of the other tRNA-LC subunits would likely equally affect RNA ligation in all tissues and cell types. As a protector against oxidative inactivation, PYROXD1 is on the other hand more critical in tissues and under the conditions of high (oxidative) stress, elevated levels of transition metal critical for RTCB oxidation or altered levels of PYROXD1 regulators NADPH and O_2_^19^. Therefore, where defects in other subunits may cause organism-wide consequences and potential lethality, severity of symptoms linked to PYROXD1 would depend on tissue type and genetic background, particularly in moderately severe variants. That is likely why, for example, the most frequent N155S variant can cause symptoms anywhere from early-onset to adulthood^30,33–35,65–69^.

Skeletal muscles, relying on high mitochondrial activity and experiencing large changes in oxygen pressure and stress levels^70^, could require a malleable and dependable protective mechanism supporting RNA ligation. Yet, why other oxidatively burdened tissues are not consistently impacted in PYROXD1 myopathy, and how this interplays with chronic stress and tissue oxygenation remains unknown. Additional symptoms do variably appear among PYROXD1 myopathy patients, ranging from cardiomyopathy^30^ to connective tissue presentation^29^, which may non-universally arise as a consequence of specific genetic background. Since PYROXD1 myopathy is a relatively newly identified disease, it remains likely under-diagnosed, with 31 patients reported so far in peer reviewed publications. Ongoing efforts are focused on identification of the full range of symptoms in all affected individuals.

Peripheral nerve involvement deserves particular attention. Neuropathic signs, predominantly hyporeflexia, are reported in roughly half of PYROXD1 patients^35^, and axonal neuropathy was documented in the oldest individuals of the original cohort^30^. This has implications for oculopharyngodistal myopathy (OPDM), caused by GGC expansions with unresolved pathomechanism^71^. Similar GGC expansions produce polyglycine that sequesters the tRNA-LC and impairs tRNA ligation in neurodegenerative disorders such as FXTAS^26^. Our demonstration that impaired RNA ligation suffices to cause a myopathy may thus not only reveal pathomechanism of PYROXD1 myopathies, but predict the basis for previously poorly understood OPDM.

It remains unknown whether tRNA-LC in skeletal muscles is particularly exposed to stress and thus in need of more robust protection, or whether skeletal muscles are instead more burdened by the loss of RNA ligation and its downstream consequences including mitochondrial deregulation. These hypotheses are not mutually exclusive. Changes in iron, oxygen, NADPH or oxidative stress levels may cause chronic alterations in tRNA-LC activity. In turn, loss of RNA ligation can remodel mitochondrial composition, together establishing a potential regulatory feedback loop. Such feedback would position tRNA-LC inactivation as a trigger for compensatory responses including mitochondrial remodelling, suggesting a regulatory link between RNA ligation and mitochondrial homeostasis with potential relevance beyond PYROXD1 myopathies. Whether this mechanism also operates under normal physiological conditions, and whether it is mediated by controlled accumulation of unligated tRNA fragments as signalling intermediates, remains to be tested.

## Acknowledgements

The authors thank the patients and families for their generous donations of biospecimens used to derive cell lines used in this research. We thank Dr. Hernan Gonorazky and Dr. Zahra Aleisa (Sick Kids Toronto, Canada), Professor Carsten Bönnemann (NIH, now St. Jude Children’s Research Hospital, USA) and Professor Henna Tyynismaa (University of Helsinki, Finland) for sharing patient fibroblast cell lines. We thank Martin Windberger for technical assistance. We would also like to thank WeiQiang Chen, Dorothea Anrather and Natascha Hartl from the Max Perutz Labs Mass Spectrometry Facility for mass spectrometry sample processing, data acquisition and analysis. Furthermore, we thank Thomas Sauer, Endre Kiss and Kitti Csályi from the Bio-optics facility from the Max Perutz Labs for FACs samples processing and assistance. We thank Stefan Schüchner and Alexander Stadler for production of antibodies used in this work, and for constructive consultations. Finally, we thank Professor Martin Jinek and Moritz Pfleiderer (University of Zurich, Switzerland) for fruitful discussions.

## Ethical statement

Patients’ samples were collected, analysed, and published with informed written consent.

## Materials and methods

### Variant nomenclature

This manuscript primarily focuses on the six single amino acid variants found in patients so far. We refer to these, and not frameshift or exon skipping pathogenic variants, unless stated otherwise. Furthermore, as the manuscript focuses on variant effects on protein structure, we use “PYROXD1 variants” throughout the manuscript, rather than alternating between the protein name and the gene name (*PYROXD1*). Similarly, we will use single letter codes, such as N155S or ΔE496, without the “p” prefix. Here, we provide the ClinVar identifiers of the single amino acid variants discussed in this manuscript:

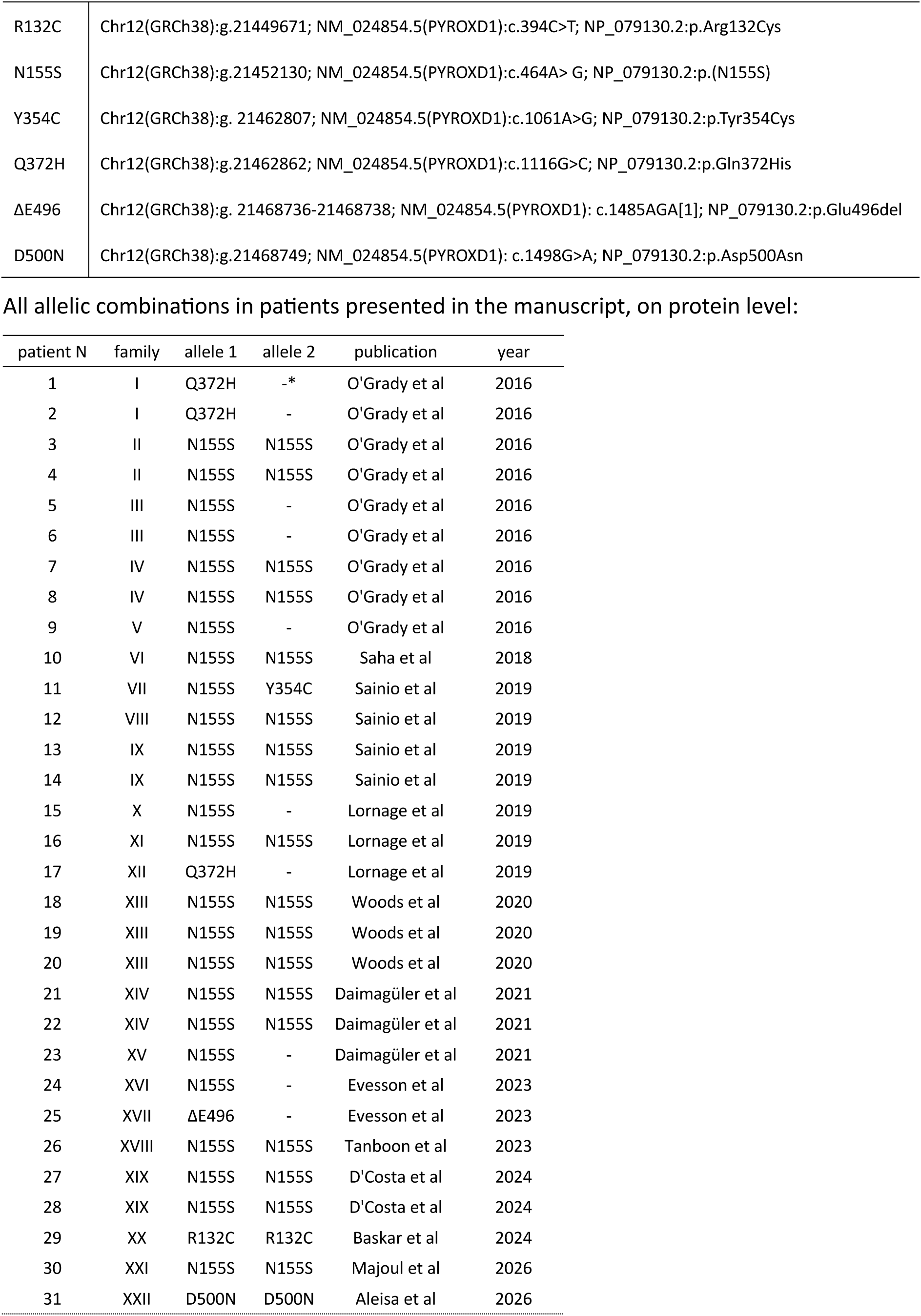
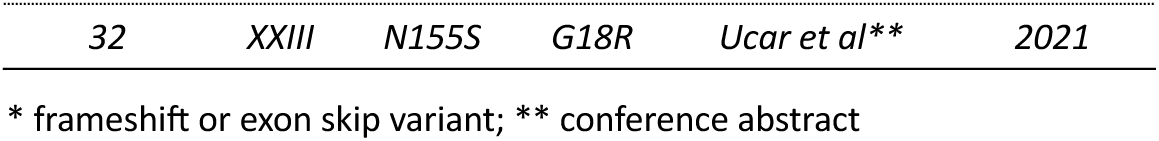

### Cell lines and culture conditions

Tet-ON HeLa cell lines expressing a Tet-regulated shRNA^9,19^ and Flp-In-T-Rex-HEK293 were cultured at 37°C, 5% CO2 in DMEM supplemented with 10% fetal calf serum, 100U·mL^-1^ penicillin, 100 μg·mL^-1^ streptomycin and 200mM HEPES. Cells were regularly tested for mycoplasma contamination.

Human fibroblasts from healthy controls and PYROXD1 patients were maintained in DMEM:F12 supplemented with 10% fetal calf serum, 5% Amniomax II complete medium, 100U·mL^-1^ penicillin and 100 μg·mL^-1^ streptomycin as described previously^29^.

### Stable cell lines generation

HEK Flp-In T-REx 293 cells overexpressing FLAG-HA-PYROXD1 variants were generated as described before^19^. In short, cells were co-transfected with the pOG44 vector and a pcDNA5/FRT/TO vector carrying the construct to be integrated under a Tet-On-controlled CMV promotor. As negative control for the selection, only the pOG44 vector was transfected. Two days after the transfection, the cells were split into selection medium [complete DMEM with 50 μg·mL^-1^ Hygromycin B]. The selection was stopped after all cells in the negative control were dead. The selected cells were then tested for construct expression upon doxycycline addition and further passaged as usual.

To generate TET-ON HeLa cells endogenously FKBP12^F36V^-tagged, we selected gRNA sequences that cut in the region surrounding the start codon of the targeted gene (TGAATATCGCGGCGGAGCAA for PYROXD1; TTCCACAAACGCACCGGAGA for DDX1) and cloned them into the pX330-U6-Chimeric_BB-CBh-hSpCas9-hGem(1/110)-P2A-mCherry vector (modified from^72^) as described previously^73^. Repair templates were cloned so that they have the following structure: left homology arm – knock-in sequence – right homology arm. The knock-in sequence either encoded BFP-P2A-FKBP12^F36^-linker (FKBP12^F36V^ knock-in on PYROXD1) or FKBP12^F36^-linker (FKBP12^F36V^ knock-in on DDX1). The homology arms contained the gDNA sequences surrounding the knock-in site (∼500 bp on each side), with the 3’ end of the left homology arm ending just after the start codon of the endogenous locus and the 5’ end of the right homology arm beginning with the first codon after the start codon of the endogenous locus. Cells were co-transfected with the corresponding repair template and the sgRNA/Cas9 vector. Two days after the transfection, the cells were sorted by FACS (BD FACSMelody) for mCherry expression. The mCherry-positive cells were expanded for 5 days and then single-cell sorted for either positive BFP and negative mCherry expression (FKBP12^F36V^ knock-in on PYROXD1) or negative mCherry expression (FKBP12^F36V^ knock-in on DDX1). The clonal cell lines were expanded for two weeks and then genotyped by PCR. Clones that had the intended homozygous knock-in were subsequently validated by Sanger sequencing of the insertion locus and by western blotting.

To generate FKBP12^F36V^-PYROXD1 Tet-ON HeLa cells stably overexpressing N-terminal FLAG-HA-PYROXD1 WT and disease variants, cells were transduced with a retroviral pBMN-1-GFP mammalian vector with addition of 5 μg·mL^-1^ polybrene. One week after transduction, the cells were sorted by FACS (BD FACSAria III) for positive GFP expression taking only the lowest 10% expressing population. Afterwards, the cells were expanded for 2 days and re-sorted by FACS (BD FACSAria III) taking the lowest 5% GFP expressing cells from the previous population. The cell lines were validated by western blot.

### Cell treatments

Tet-ON HeLa cells were treated with 2 μg·mL^-1^ doxycycline for 7 days following the next regime: Day 0 – seed and treated; day 2 – doxycycline treatment; day 4 – split and treated; day 6 – doxycycline treatment; day 7 – harvest for downstream analysis.

HEK293 cells were treated with 2 μg·mL^-1^ doxycycline for 24 hours and harvested for downstream analysis.

Endogenously FKBP12-PYROXD1 tagged Tet-ON HeLa cell lines were treated with 250nM dTAG^V-1^ or dTAG^V-1NEG^ as a negative control for 3 days. The degradation drug was added every day to the cells. For viability assays, cells were treated with 250nM dTAG^V-1^ or dTAG^V-1NEG^ for 3 days, split and re-treated for 2 more days.

For UPR induction, cells were treated with 1.5 μg·mL^-1^ tunicamycin for 4 h.

### Transient transfections

Cells were seeded on 6-well plates (400,000 cells per well) and treated with 250nM dTAG^V-1^ overnight. The next day, cells were treated with 125nM dTAG^V-1^ and transfected with 1 μg of pcDNA5/FRT vector lacking the Tet operator expressing FLAG-HA-PYROXD1 using Lipofectamine 3000 according to the manufacturer’s recommendations. 6 hours later, the medium was changed to DMEM containing 250nM of dTAG^V-1^. 2 days after transfection, the cells were harvested for downstream analysis.

### Construct cloning and mutagenesis

Expression constructs were cloned into the pcDNA5/FRT/TO Tet-On-controlled CMV promoter for HEK293 stable cell line generation, pBMNI-1-GFP for Tet-ON HeLa stable cell line generation and into pcDNA5 with the CMV promoter without the Tet-On-system for transient transfection. pGEX-6P-1 vector was used for recombinant protein expression. Constructs were assembled from gel-purified PCR products or from synthesized DNA fragments using NEBuilder HiFi DNA Assembly Master Mix (NEB, E262), according to the manufacturer’s instructions. PCR fragments were amplified using Q5 Hot Start High-Fidelity DNA Polymerase (NEB, M0493), either using cDNA or existing lab-internal plasmids as template. Primers for site-directed mutagenesis of constructs were designed with NEBaseChanger (https://nebasechanger.neb.com/) and the linear mutagenized constructs were generated with Q5 Hot Start High-Fidelity DNA Polymerase (NEB, M0493) from the non-mutagenized constructs. For circularization of linear mutagenized constructs, 2 μL of the unpurified PCR product was mixed with 2 μL 10x rCutSmart (NEB, B6004), 2 μL 10 mM ATP, 1 μL 100 mM DTT, 1 μL T4 DNA Ligase (NEB, M0202), 0.2 μL T4 PNK (NEB, M0201), 0.2 μL DpnI (NEB, R0176) and 11.6 μL nuclease-free water. The circularization reaction was performed in a thermocycler: 15 min at 37 °C, followed by 5 min at 23 °C. Cloning reactions, both from general cloning and mutagenesis, were transformed into competent NEB 5-alpha E. coli cells (NEB, C2987) and resulting colonies were further grown as liquid cultures, from which plasmid DNA was isolated using Qiagen mini kit. All generated constructs were validated by Sanger sequencing.

### Expression and purification of recombinant PYROXD1

Recombinant GST-tagged PYROXD1 WT and mutant proteins were expressed and purified essentially as previously described^14,21^. Briefly, proteins were expressed in *E. coli BL21(DE3)*, purified with glutathione Sepharose 4B beads, subjected to GST-tag cleavage, and subjected to two rounds of gel filtration in a HiLoad Superdex 75 16/60 column. Purified proteins were concentrated to 40 μM, flash frozen and stored at -80°C until use.

### Expression and purification of recombinant tRNA ligase complex

Recombinant tRNA ligase complex was expressed and purified as previously described^15,19^. In short, insect cells were resuspended in lysis buffer (20 mM HEPES pH 8.0, 150 mM NaCl, 0.1 % Tween-20, cOmplete Protease Inhibitor Cocktail (Roche)), and lysed by sonication. The supernatant was applied to a Ni-NTA Super-flow resin (QIAGEN) and eluted with buffer containing 20 mM HEPES pH 8.0, 500 mM NaCl, 0.5 mM TCEP, and 250 mM imidazole. The fusion tags were removed by proteolysis using His6-tagged Tobacco Etch Virus (TEV) protease during an overnight dialysis at 4°C against 20 mM HEPES pH 8.0 and 150 mM NaCl. The protein solution was subsequently supplemented with 0.5 mM TCEP and applied to a Ni-NTA Superflow resin (QIAGEN) to remove any uncleaved protein, His6-TEV protease, and remaining impurities. The complex-containing flow-through was loaded onto a size-exclusion chromatography column (Superdex 200, GE Healthcare) and eluted in buffer containing 20 mM HEPES pH 8.0, 150 mM KCl, and 0.5 mM TCEP. Peak fractions containing the tRNA ligase complex were concentrated to 9.4 mg·ml-1 using a centrifugal filter (Amicon Ultra, MWCO 30 kDa, Sigma), flash frozen in liquid nitrogen and stored at -80°C.

### Thermal shift assay

Melting temperatures and Tm values of PYROXD1 variants were assessed by fluorescence-based thermal shift assay. Recombinant PYROXD1 [10μM] was mixed in buffer containing 0.75mM HEPES pH 7.4, 2.25mM KCl, 0.75mM MgCl₂ and 104 μM AEBSF with SYPRO Orange at a final concentration of 0.125×. Samples were prepared in 96-well plates in at least three technical replicates. The fluorescence signal was monitored upon temperature increase from 10 to 95 °C, at a heating rate of 1.5 °C/min using Bio-Rad CFX96. To account for curve asymmetry, in particular in presence of NADPH, melting curve derivative centroids were used as melting point estimates.

### Steady state measurements

Equal volumes (100 μL) of solution of NADPH (solution 1) and a solution of recombinant PYROXD1 and the electron acceptor substrate (100 μL) (solution 2) were mixed and absorbance at 340 nm was measured using Synergy H1 Hybrid Multi-Mode Microplate Reader at room temperature. The assay buffer was 0.75mM HEPES pH 7.4, 2.25mM KCl, 0.75mM MgCl₂ and 104 μM AEBSF. The final concentration of NADPH was 350 μM NADPH. The final concentration of PYROXD1 was 20 μg·L^-1^ in all experiments. The final concentrations of electron acceptors were: menadione 70 μM, ethanol (as O_2_ donor). Volume of the added stock solution never exceeded 2.5% of the reaction volume. Reaction mixtures without enzyme (enzyme replaced with buffer) and without menadione (ethanol) were used as negative controls. Enzymatic activities were measured based on the slope of the absorbance decrease [min^-1^], and using optical path 0.56 cm, reaction volume 0.2 mL and extinction co-efficient 6220 M^−1^·cm^−1^ to calculate the specific activity. One unit of activity was defined as the amount of enzyme that oxidizes 1 mM of NADPH in 1 min. All measurements were done at room temperature. Significance of average values for all samples was evaluated by comparing values obtained for the sample measurements with the values obtained for the corresponding control using t test assuming unequal variances, s = 0.05.

### tRNA splicing assay

The assay was performed as previously described^19^. Briefly, one volume of 0.1 mM body-labeled S. cerevisiae pre-tRNA-Phe^GAA^, pre-heated at 95°C for 60s and incubated for 20 min at room temperature, was mixed with four volumes of reaction buffer (100 mM KCl, 5.75 mM MgCl2, 2.5 mM DTT, 5 mMATP, 6.1 mM Spermidine-HCl pH 8.0 (Sigma), 100 U·mL^-1^ RNasin RNase inhibitor [Promega]). Equal volumes of this reaction mixture and cell extracts (protein concentration 3-6 mg·mL^-1^) were mixed and incubated at 30°C for 30 minutes. The reaction mix was sub-sequently deproteinized with proteinase K, followed by phenol/chloroform extraction and ethanol precipitation. Reaction products were separated on a 10% denaturing polyacrylamide gel, and mature tRNA formation and tRNA exon formation was monitored by phosphorimaging.

### RNA ligation assays

Substrate preparation for intra-strand and inter-strand ligation assays was carried out as previously described^19^. In short, 50 pmol RNA oligonucleotide derived from the firefly luciferase gene (5’-UCGAAGUAUUCCGCGUACGU-3’, Dharmacon) were incubated with 1.11 MBq [5’-32P] cytidine-3’,5’-bisphosphate (111 TBq·mmol^-1^, Perkin Elmer) and 20 units T4 RNA ligase 1 (NEB) for 1 hr at 16 C. Labeling reactions were resolved by denaturing gel electrophoresis in 15% poly-acrylamide gels containing 8 M urea (SequaGel, National Diagnostics). The 3’ end-labeled RNA was visualized by autoradiography and passively eluted from gel slices overnight at 4 C in 0.3 M NaCl.

#### Intra-strand ligation (circularization) assay (for recombinant proteins)

Activity of the recombinant tRNA-LC and RTCB was measured using an RNA (circularization) ligation assay. In short, the solution containing 12.5 mM TCEP, 100 mM KCl, 2.9 mM MgCl2, 7.5 mM ATP, 0.5 mM GTP, 250 mM ZnCl_2_, Rnasin Ribonuclease Inhibitors (1 mL per 500 mL) and 65% (v/v) glycerol was first mixed (2/1, v/v) with radiolabelled oligoribonucleotide to prepare the reaction cocktail. The cocktail was then mixed (3/2, v/v) with the sample containing recombinant tRNA-LC or RCTB, and Archease, and incubated for 30 min at 30 C. The reaction was then stopped with 2xFA solution (1/1, v/v), composed of 90% Formamide, 50 mM EDTA, 1 ng·mL^-1^ bromophenol blue, 1ng·mL^-1^ Xylene Cyanol and boiled for 1 min. The reaction products were separated on a denaturing 15% urea-polyacrylamide gel (SequaGel) and detected by autoradiography. ImageQuant software was used to quantify the band intensities and substrate conversion rate was calculated as the quotient of the substrate and the sum of substrate and product band intensities. When done in independent replicates, the results are represented as the average value with the standard error of the mean (s = 0.05).

#### Inter-strand ligation assay (for cell extracts)

For the use in interstrand ligation assays using cell extracts, the RNA was annealed to a non-labeled, complementary RNA oligonucleotide (5’-CGUACGCGGAAUACUUCGA-3’, Dharmacon, Thermo Scientific). 3/5th volumes of a reaction mixture (67 mM KCl, 2 mM MgCl2, 8.3 mM DTT, 5 mM ATP, 0.3 mM GTP, 53 U·mL^-1^ RNasin RNase inhibitor, 43% glycerol) containing 17 nM radiolabelled RNA duplex were mixed with 2/5 volumes of cell extracts (protein concentration 3-6 mg·mL^-1^) and incubated at 30°C for 30 minutes. Reaction products were subsequently separated on a 15% denaturing polyacrylamide gel, and ligation products were monitored by phosphorimaging. Quantification of band intensities was performed using the ImageQuant software.

### In vitro reconstitution experiment

Recombinant PYROXD1 (100nM), tRNA-LC (4 subunits, untruncated) (25 nM) and Archease (200 nM) were pre-incubated with 600 μM H_2_O_2_ in a buffer containing 30 mM HEPES/KOH pH 7.4, 100 mM KCl, 5 mM MgCl2, 104 μM AEBSF, 10% glycerol, and 1% NP40. Afterwards, the pre-incubation was stopped by a reaction cocktail containing TCEP, ATP, GTP and ssRNA oligonucleotide (See RNA ligation assays; 2/3, v/v) and the activity of the tRNA-LC was measured using the intra-strand RNA ligation (circularization) assay.

### UV-Visible Spectroscopy

Frozen aliquots of recombinant PYROXD1 were thawed and immediately put at 4°C, span down at 13000rpm for 2 mins. UV-visible measurements were performed in technical triplicates by nanodrop using a DeNovix DS-11+ Spectrophotometer at room temperature.

### SDS-PAGE of recombinant PYROXD1

Frozen aliquots of recombinant PYROXD1 were thawed and immediately put at 4°C, span down at 13000rpm for 2 mins and separated by SDS-PAGE on 10% polyacrylamide gels. Gels were stained in InstantBlue Coomassie Protein Stain (ISB1L), subsequently destained in deionized water and imaged using a ChemiDoc imaging system.

### In vitro immunoprecipitation and time course

Recombinant FLAG-HA-PYROXD1 and variants (100 nM) were incubated with recombinant tRNA ligase complex (180 nM) in 100 µL of buffer [30 mM HEPES/KOH pH 7.4, 100 mM KCl, 5 mM MgCl2, 104 μM AEBSF, 10% glycerol, 1% NP40] and 1 mM NADPH for 20 min on ice. Following incubation, 12.5 µL of anti-FLAG M2 Affinity gel equilibrated in buffer were added to each reaction and samples were incubated for 90 minutes at 4°C with gentle rotation. The beads were subsequently collected by centrifugation in a picofuge and washed 4 times with 100 µL NADPH-containing buffer [0.5mM].

Bound proteins were eluted by incubation with 13µl 3xFLAG-peptide for 60 min at 4°C while shaking. Eluted samples were analyzed by SDS-PAGE followed by silver staining.

### Co-immunoprecipitation

HEK293 cells expressing FLAG-HA-PYROXD1 (see stable cell lines generation) and a control, untagged cell line were treated with doxycycline (see cell treatments). Cells were then harvested and lysed in lysis buffer [30 mM HEPES/KOH pH 7.4, 100 mM KCl, 5 mM MgCl2, 104 μM AEBSF, 10% glycerol, 1% NP40] and incubated with 1 mM NADPH or no nucleotide. After taking 2.5% of the lysate as Input, Anti-FLAG M2 Affinity Gel (Sigma) was added to the extracts and incubated on a rotating wheel at 4C for 1h for FLAG-PYROXD1 immunoprecipitation. For supernatant experiments, 2.5% of the unbound fraction was taken after incubation. Beads were then washed five times with the same buffer. Bound complexes were eluted in 0.5 mg·mL^-1^ 3xFLAG peptide dissolved in lysis buffer for 45 min, 4°C, while shaking. 5x SDS loading dye (5 % SDS w/ v, 62.5 mM Tris pH 6.8, 50 % Glycerol v/v, 25 mM EDTA, 0.025 % Bromophenol blue w/v, 5% β-Mercaptoethanol v/v) was added to the eluates, the input samples, as well as the supernatant samples. The samples were mixed, boiled at 98 °C for 1 min and then either sent to Mass Spectrometry or were analysed by western blot.

### RT-qPCR

cDNA was generated from 1 μg of total RNA using the Maxima First Stand cDNA synthesis kit, following manufacturer’s instructions, and was diluted 1:6 in nuclease-free water before downstream analysis. For RT-qPCR, a reaction containing 2 μl of cDNA, 5 μl 2x GoTaq (Promega), 0.8 μl primer mix (5μM each) and 2.2 μl nuclease-free water was prepared in a well of a 384-well plate. The RT-qPCR was performed in a CFX384 Touch thermocycler (Bio-Rad) using the following protocol: 95°C for 5 min, 95°C for 10s, 60°C for 30s, plate read, repeat previous steps 50 cycles, 95°C for 10s, melt curve determination from 55°C to 95°C in 0.5°C increments. The quality of the PCR products was evaluated by melting curve analysis. Relative expression levels were calculated from Ct values with the 2^-ΔΔCt^ method, normalizing expression levels to *ACTB* mRNA levels (Livak & Schmittgen, 2001b)

### Western blotting

Cells were harvested and lysed in lysis [30 mM HEPES pH 7.4, 100 mM KCl, 5 mM MgCl₂, 104 μM AEBSF, 10% (v/v) glycerol and 1% (v/v) NP-40]. Protein concentrations were determined using the Bradford assay, and normalized to equal amounts of protein. The samples were mixed with 5xSDS loading dye and separated by SDS-PAGE on 10% polyacrylamide gels. Proteins were transferred onto Immuno-Blot PVDF membranes (Bio-Rad) using a semi-dry transfer system. Membranes were blocked with 5% non-fat milk in PBS-T for 1 hour at room temperature and subsequently incubated overnight at 4°C with primary antibodies against PYROXD1, RTCB, Archease, FAM98B, DDX1, CGI-99 and GAPDH. The membranes were washed 3 times with PBS-T and incubated with HRP-conjugated secondary antibodies for 1 h at room temperature. Afterwards, the membranes were washed 3 times with PBS-T and signals were detected using Clarity Western ECL Substrate (Bio-Rad) using a ChemiDoc imaging system. For quantification, band intensities were measured using Image Lab. Data were obtained from independent biological replicates and unpaired t-test was used for statistical analysis.

### Northern blotting

Total RNA was isolated from HeLa cells in 6-well plates and human fibroblasts in 6cm plates using TRIzol, following the manufacturer’s instructions. 5-6 μg of RNA was mixed with equal volumes of 2X FA buffer (90% formamide, 50 mM EDTA pH 8.0, 0.025 % Bromophenol blue w/v, 0.025 % Xylene cyanol w/v) and separated in a 10% UREA-PAGE. The RNA was blotted on Hybond-N+ membrane and fixed by ultraviolet cross-linking. Membranes were pre-hybridized in 5X SSC, 20 mM Na2HPO4 pH 7.2, 7 % SDS, and 3 mg sonicated salmon sperm DNA for 1 hr at 50°C. Hybridization was performed in the same buffer overnight with 100pmol of [5’-^32^P]-labeled DNA probes. Subsequently, the blots were washed twice for 1min with 5X SSC, 5% SDS and once for 1 min with 1X SSC, 1% SDS and analyzed by phosphoimaging in an Amersham Typhoon imager.

### Silver staining

Proteins were separated by SDS-PAGE and visualized using a modified silver staining procedure based on Blum et al., 1986. Gels were fixed in methanol/acetic acid/water (500:120:380, v/v/v) for at least 30 min, washed twice with 50% ethanol for 5 min and once with 30% ethanol for 5 min, and subsequently treated with 0.8 mM sodium thiosulfate for 1 min. Following three washes with water, gels were incubated for 20 min in a solution containing 0.2 g·L^-1^ silver nitrate and 0.7 mL·L^-1^ 37% formaldehyde. After three washes with water, gels were developed in 60 g·mL^-1^ sodium carbonate containing 0.5 mL·L^-1^ 37% formaldehyde and 4 mg·L^-1^ sodium thiosulfate. Development was monitored by imaging at regular intervals using a ChemiDoc imaging system and terminated with 5% acetic acid.

### Cell viability assay

Cells were seeded on 12-well plates and treated with doxycycline or dTAG^V-1^ (see cell treatments) for 3 days. The treatment was started for each well independently depending on the number of days of treatment (1 to 5-7 days). Subsequently, the cells were harvested and seeded on 96-well plates (2,000 cells per well) and treated with the corresponding drug for 4 or 2 days more, respectively. Cell viability was assessed using AquaBluer as an indicator of viable cells according to the manufacturer’s instructions.

### Statistical Analysis

Due to high sensitivity of RTCB and PYROXD1 to slight changes in buffer conditions including trace metal ions, we always compared the outcomes of variants or treatments relative to control or WT conditions, unless designated otherwise. We first harmonized the entire datasets between biological replicates across mean value for each set, followed by normalization relative to median value of control or WT condition. Unless stated otherwise, such as in in large scale sample analysis (for example, see mass spectrometry analysis below), we used unpaired Student’s t test in GraphPad Prism 11 assuming unequal variances, with * p < 0.05; ** p < 0.01; *** p < 0.001; **** p<0.0001. Also, unless stated otherwise, experiments in cell culture were done in biological replicates, and experiments with recombinant proteins involved analysis of one aliquot per datapoint, and where designated batches of recombinant protein preparations.

## Mass Spectrometry

### Analysis of mouse hearts

#### Sample preparation for mass spectrometry analysis

Cell pellets were resuspended in 2% sodium deoxycholate (SDC), 0.1 M Tris-HCl (pH 8.8), and heated at 95 °C for 10 min with shaking (1,000 rpm). After cooling, lysates were sonicated in a Bioruptor (5 cycles, 30 s on/30 s off, high power, 4 °C cold room), treated with 0.5 µL benzonase for 15 min on ice, and sonicated again under the same conditions. Debris was removed by centrifugation (15,000 × g, 10 min, 10 °C). Proteins in the supernatant were precipitated with four volumes of ice-cold acetone (−20 °C) overnight at 4 °C, pelleted (15,000 × g, 10 min, 10 °C), washed with 500 µL of 80% acetone (−20 °C), and air-dried for 5 min. Pellets were redissolved in 100 µL of 2% SDC, 50 mM ammonium bicarbonate (ABC) by sonication as above, and protein concentration was determined by micro-BCA assay (Thermo Fisher Scientific).

For digestion, approximately 100 µg protein (20 µL) was reduced with 10 mM DTT (30 min, 50 °C) and alkylated with 20 mM iodoacetamide (30 min, room temperature, in the dark). Excess iodoacetamide was quenched with 2.5 mM DTT (10 min). Samples were diluted to 1% SDC with 50 mM ABC and digested with 2 µg trypsin (Trypsin Gold, Promega) at 37 °C overnight. Digestion was stopped with TFA (1% final), and SDC precipitate was removed by centrifugation (10,000 × g, 10 min). The supernatant was acidified further to 2% TFA, centrifuged again (10,000 × g, 10 min), and desalted on C18 StageTips^74^.

#### Liquid chromatography-mass spectrometry analysis

LC-MS analysis was performed on an UltiMate 3000 RSLCnano LC system (Thermo Scientific) coupled to a timsTOF HT (Bruker). The system was equipped with a CaptiveSpray ion source (Bruker), and a Butterfly Portfolio Heater (Phoenix S&T).

Peptides were loaded onto a trap column (PepMap Neo C18 5mm × 300 µm, 5 μm particle size, Thermo Scientific) using 0.1% TFA as mobile phase, and separated on an analytical column (Aurora Ultimate CSI C18, 25 cm × 75 µm, 1.7 µm particle size, IonOpticks), applying a linear gradient starting with a mobile phase of 98% solvent A (0.1% FA) and 2% solvent B (80% acetonitrile, 0.08% FA), increasing to 35% solvent B over 60 min at a flow rate of 300 nl/min. The analytical column was heated to 50°C.

The mass spectrometer was operated in data-independent acquisition (DIA) parallel accumulation serial fragmentation (PASEF) mode. MS2 data were acquired with eight PASEF scans per duty cycle, each containing three m/z windows. The m/z window widths were adjusted based on the expected precursor density, covering a total range of 300–1200 m/z. The ion mobility range was set to 0.64-1.42 V*s/cm, and the accumulation and ramp time was set to 100 ms. TIMS elution voltages were calibrated linearly to obtain the reduced ion mobility coefficients (1/K0) using three Agilent ESI-L Tuning Mix ions (m/z 622, 922 and 1,222). Collision energy for fragmentation was scaled linearly with precursor mobility (1/K0), ranging from 20 eV (at 1/K0 = 0.6 V*s/cm) to 59 eV at (at 1/K0 = 1.6 V*s/cm).

#### Data analysis

MS raw data were converted to htrms format using HTRMS converter (Biognosys). Then it was searched with Spectronaut (19, Biognosys) in directDIA+ mode against the Uniprot mouse reference proteome (version 2024_01, www.uniprot.org), as well as a database of most common contaminants (release 2023_03, https://github.com/maxperutzlabs-ms/perutz-ms-contaminants). The search was performed with full trypsin specificity and a maximum of two missed cleavages at a protein and peptide spectrum match false discovery rate of 1%. Carbamidomethylation of cysteine residues was set as fixed, oxidation of methionine, and N-terminal acetylation as variable modifications. The cross-run normalization was turned off and all other settings were left as default.

Computational analysis was performed using Python and the Python library MsReport (version 0.0.24; source code: https://github.com/hollenstein/msreport).

Only non-contaminant proteins identified with a minimum of two peptides and quantified in at least four replicates of one condition were considered for further analysis. LFQ protein intensities were log2-transformed and normalized across samples using mode normalization from MsReport, which involves calculating log2 protein ratios for all pairs of samples and determining normalization factors based on the modes of the resulting ratio distributions. Missing values were imputed by drawing random values from a normal distribution. The parameters of this distribution were calculated per sample from the standard deviation and median of the observed log2 protein intensities (μ = median sample LFQ intensity – 1.8 standard deviations of the sample LFQ intensities, σ = 0.3 × standard deviation of the sample LFQ intensities). iBAQ intensities were calculated by dividing protein intensities by the number of theoretically observable tryptic peptides between 7 and 30 amino acids. Statistical analysis was performed using the linear models for microarray analysis (limma) v.3.54.2 package in R. Moderated *t*-statistics were calculated using the limma-trend method. Multiple testing correction was applied using the Benjamini–Hochberg method.

### Analysis of immunoprecipitates

#### Sample preparation for mass spectrometry analysis

Protein samples (14 µL) were mixed with 50 µg of a 1:1 mixture of Sera-Mag SpeedBeads (GE Life Sciences, cat. nos. 45152105050350 and 65152105050350) in 1 µL, and 15 µL ethanol was added to reach a final ethanol concentration of 50%. Samples were incubated in a thermomixer for 15 min at 24 °C with shaking (1,000 rpm) to allow protein binding. The supernatant was removed on a magnetic rack, and the beads were washed four times with 180 µL of 80% ethanol in water, off the rack. For on-bead digestion, 40 µL of 1.25 mM TCEP and 9.6 µL of 5 mM iodoacetamide in 50 mM ammonium bicarbonate (ABC) were added together with 300 ng trypsin (Trypsin Gold, Promega) in 3 µL of 50 mM ABC, and samples were incubated at 37 °C overnight with shaking (1,000 rpm), allowing reduction, alkylation, and digestion to proceed simultaneously. The beads were collected on a magnetic rack, and the supernatant was transferred to a new tube. The beads were rinsed with 20 µL of 50 mM ABC and sonicated for 30 s, and the resulting supernatant was pooled with the first. Digests were acidified with trifluoroacetic acid (TFA) to a final concentration of 0.5%.

#### Liquid chromatography-mass spectrometry analysis

LC-MS analysis was performed on a Vanquish Neo UHPLC system (Thermo Scientific) coupled to a timsTOF HT (Bruker). The system was equipped with a CaptiveSpray ion source (Bruker), and a Butterfly Portfolio Heater (Phoenix S&T).

Peptides were loaded onto a trap column (PepMap Neo C18 5mm × 300 µm, 5 μm particle size, Thermo Scientific) using 0.1% TFA as mobile phase, and separated on an analytical column (Aurora Ultimate CSI C18, 25 cm × 75 µm, 1.7 µm particle size, IonOpticks), applying a linear gradient starting with a mobile phase of 98% solvent A (0.1% FA) and 2% solvent B (80% acetonitrile, 0.08% FA), increasing to 35% solvent B over 60 min at a flow rate of 300 nl/min. The analytical column was heated to 50°C.

The mass spectrometer was operated in data-independent acquisition (DIA) parallel accumulation serial fragmentation (PASEF) mode. MS2 data were acquired with eight PASEF scans per duty cycle, each containing three m/z windows. The m/z window widths were adjusted based on the expected precursor density, covering a total range of 300–1200 m/z. The ion mobility range was set to 0.64-1.42 V*s/cm, and the accumulation and ramp time was set to 100 ms. TIMS elution voltages were calibrated linearly to obtain the reduced ion mobility coefficients (1/K0) using three Agilent ESI-L Tuning Mix ions (m/z 622, 922 and 1,222). Collision energy for fragmentation was scaled linearly with precursor mobility (1/K0), ranging from 20 eV (at 1/K0 = 0.6 V*s/cm) to 59 eV at (at 1/K0 = 1.6 V*s/cm).

#### Data analysis

MS raw data were converted to htrms format using HTRMS converter (Biognosys). Then it was searched with Spectronaut (21.0, Biognosys) in directDIA+ mode against the Uniprot human reference proteome (version 2026_01, www.uniprot.org), as well as a database of most common contaminants (release 2026_01, https://github.com/maxperutzlabs-ms/perutz-ms-contaminants). The search was performed with full trypsin specificity and a maximum of two missed cleavages at a protein and peptide spectrum match false discovery rate of 1%. Carbamidomethylation of cysteine residues was set as fixed, oxidation of methionine, and N-terminal acetylation as variable modifications. The cross-run normalization was turned off and all other settings were left as default.

Computational analysis was performed using Python and the Python library MsReport (version 0.0.34; source code: https://github.com/hollenstein/msreport).

Only non-contaminant proteins identified with a minimum of two peptides and quantified in at least two replicates of one condition were considered for further analysis. LFQ protein intensities were log2-transformed and normalized across samples using mode normalization from MsReport, which involves calculating log2 protein ratios for all pairs of samples and determining normalization factors based on the modes of the resulting ratio distributions. Missing values were imputed by drawing random values from a normal distribution. The parameters of this distribution were calculated per sample from the standard deviation and median of the observed log2 protein intensities (μ = median sample LFQ intensity – 1.8 standard deviations of the sample LFQ intensities, σ = 0.3 × standard deviation of the sample LFQ intensities). iBAQ intensities were calculated by dividing protein intensities by the number of theoretically observable tryptic peptides between 7 and 30 amino acids. Statistical analysis was performed using the linear models for microarray analysis (limma) v.3.54.2^75,76^ package in R. Moderated *t*-statistics were calculated using the limma-trend method, including batch correction with replicate as the batch covariate. Multiple testing correction was applied using the Benjamini–Hochberg method.

### Whole cell proteome analysis

#### Sample preparation for mass spectrometry analysis

Cell pellets were resuspended in 2% sodium deoxycholate (SDC), 0.1 M Tris-HCl (pH 8.8), preheated, and incubated at 95 °C for 10 min with shaking (1,000 rpm). After cooling, lysates were sonicated in a Bioruptor (5 cycles, 30 s on/30 s off, high power, 4 °C cold room), treated with 0.5 µL benzonase for 15 min on ice, and sonicated again under the same conditions. Cell debris was removed by centrifugation (15,000 × g, 10 min, 10 °C), and the supernatant was transferred to a new tube. Protein concentration was determined by micro-BCA assay (Thermo Fisher Scientific).

For digestion, approximately 50 µg protein (25 µL) was reduced with 1 µL of 250 mM dithiothreitol (DTT; ∼10 mM final) for 30 min at 50 °C and alkylated with 1 µL of 500 mM iodoacetamide (∼19 mM final) for 30 min at room temperature in the dark. Excess iodoacetamide was quenched with 0.5 µL of 250 mM DTT for 10 min. Samples were diluted to 1% SDC with 25 µL of 50 mM ammonium bicarbonate (ABC) and digested with 0.5 µg LysC (mass spectrometry grade, FUJIFILM Wako) in 0.5 µL of 50 mM ABC for 3 h at room temperature, followed by 1 µg trypsin (Trypsin Gold, Promega) in 1 µL of 50 mM ABC at 37 °C overnight. Digestion was stopped by adding TFA to a final concentration of 2%. Samples were centrifuged (10,000 × g, 10 min), and the supernatant was transferred to a new tube and desalted on C18 StageTips^74^.

### Liquid chromatography-mass spectrometry analysis

LC-MS analysis was performed on a Vanquish Neo UHPLC system (Thermo Scientific) coupled to an Orbitrap Astral mass spectrometer (Thermo Scientific). The system was equipped interface (Thermo Scientific), a Nanospray Flex ion source (Thermo Scientific), and a Column Heater (IonOpticks) connected to a Heater Controller (IonOpticks)

Peptides were loaded onto a trap column (PepMap Neo C18 5mm × 300 µm, 5 μm particle size, Thermo Scientific) using 0.1% TFA as mobile phase, and separated on an analytical column (Aurora Ultimate XT C18, 25 cm × 75 µm, 1.7 µm particle size, IonOpticks), applying a linear gradient starting with a mobile phase of 98% solvent A (0.1% FA) and 2% solvent B (80% acetonitrile, 0.08% FA), increasing to 35% solvent B over 60 min at a flow rate of 300 nl/min. The analytical column was heated to 50°C.

The mass spectrometer Orbitrap Astral was operated in data-independent acquisition (DIA) mode. Survey scans were acquired from 380-980 m/z, normalized AGC target of 300%, resolution of 240,000 in the orbitrap with a maximum 5ms injection time. In total, 150 MS2 spectra were acquired across a mass range of 380.4-980.7 m/z using an isolation window of 4 m/z with 0 m/z overlap between windows. Window placement optimization is on. Selected ions were analyzed using a maximum injection time of 5ms, normalized AGC target of 500% after HCD fragmentation with normalized collision energy of 25% in the Astral detector. MS2 scan range is 150-2000. The loop control is time with a 0.6s cycle.

## Data analysis

MS raw data were converted to htrms format using HTRMS converter (Biognosys). Then it was searched with Spectronaut (21.1, Biognosys) in directDIA+ mode against the Uniprot human reference proteome (version 2026_01, www.uniprot.org), as well as a database of most common contaminants (release 2026_01, https://github.com/maxperutzlabs-ms/perutz-ms-contaminants). The search was performed with full trypsin specificity and a maximum of two missed cleavages at a protein and peptide spectrum match false discovery rate of 1%. Carbamidomethylation of cysteine residues was set as fixed, oxidation of methionine, and N-terminal acetylation as variable modifications. The cross-run normalization was turned off and all other settings were left as default.

Computational analysis was performed using Python and the Python library MsReport (version 0.0.34; source code: https://github.com/hollenstein/msreport)^76^.

Only non-contaminant proteins identified with a minimum of two peptides and quantified in at least two replicates of one condition were considered for further analysis. LFQ protein intensities were log2-transformed and normalized across samples using mode normalization from MsReport, which involves calculating log2 protein ratios for all pairs of samples and determining normalization factors based on the modes of the resulting ratio distributions. Missing values were imputed by drawing random values from a normal distribution. The parameters of this distribution were calculated per sample from the standard deviation and median of the observed log2 protein intensities (μ = median sample LFQ intensity – 1.8 standard deviations of the sample LFQ intensities, σ = 0.3 × standard deviation of the sample LFQ intensities). iBAQ intensities were calculated by dividing protein intensities by the number of theoretically observable tryptic peptides between 7 and 30 amino acids. Statistical analysis was performed using the linear models for microarray analysis (limma) v.3.54.2^75^ package in R. Moderated *t*-statistics were calculated using the limma-trend method. Multiple testing correction was applied using the Benjamini–Hochberg method.

