## Supplementary Figures 1-5 for "Myopathy-causing variants of PYROXD1 converge on impaired RNA ligation"

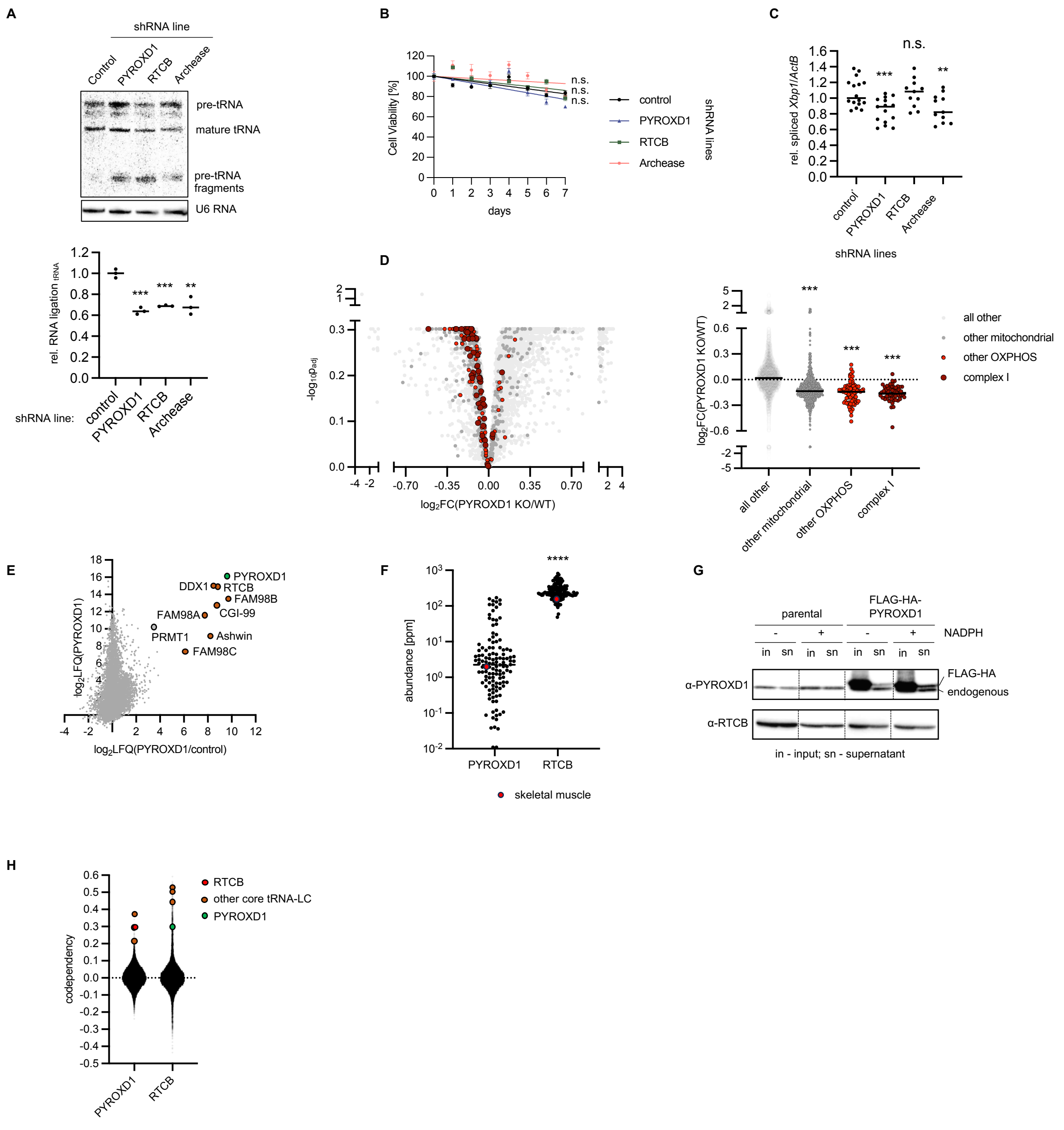


**Figure S1 (related to Figure 1). A**. PYROXD1, RTCB and Archease were silenced for 7 days in HeLa cells using genetically encoded, doxycycline-induced shRNA, and pre-tRNA fragment ligation ($Tyr^{GTA}$) was assessed by northern blotting and quantified relative to control cell line. The experiment was conducted 3 times. *** p<0.001. ** p<0.01 (student t-test) **B**. PYROXD1, RTCB and Archease were silenced as in (A) and cellular viability was monitored using AquaBluer staining. N=6. Extra sum-of-squares F test was used for the whole curve comparison. **C**. PYROXD1, RTCB and Archease were silenced as in (A), and the cells were treated with tunicamycin for 3.5 h. *XBP1* and *Actin* mRNA levels were estimated using RT-qPCR and quantified relative to control cell line. Experiment was conducted 11 times. *** p<0.001 ** p<0.01 (student t-test). **D**. PYROXD1 was inducibly and conditionally knocked-out from adult mouse hearts, which were harvested three weeks upon tamoxifen injections and ground in liquid nitrogen using freezer-mill. Protein content of the hearts was analysed using mass spectrometry. Five mice were used per condition. Mitochondrial proteins were identified with MitoCarta3.0 database. *** p<0.001 (student t-test). **E**. FLAG-HA-PYROXD1 was immunoprecipitated from HEK293 Flp-In cells with NADPH spiked into lysis and washing buffers, and the protein content in the immunoprecipitate was analysed by mass spectrometry. N=4. Subunits of the tRNA-LC and the associated protein PRMT1 are labelled. **F**. Abundance data for PYROXD1 and RTCB were obtained from paxdb6.1 database across all available tissues. Each dot represents protein abundance in the corresponding sample. **** p<0.0001 (student t-test) **G**. FLAG-HA-PYROXD1 was immunoprecipitated from HEK293 Flp-In cells with NADPH spiked in where indicated, and PYROXD1 and RTCB levels were estimated by western blotting. **H**. Cellular co-dependency data were obtained from depmap portal (26Q1). Each dot represents a single gene.

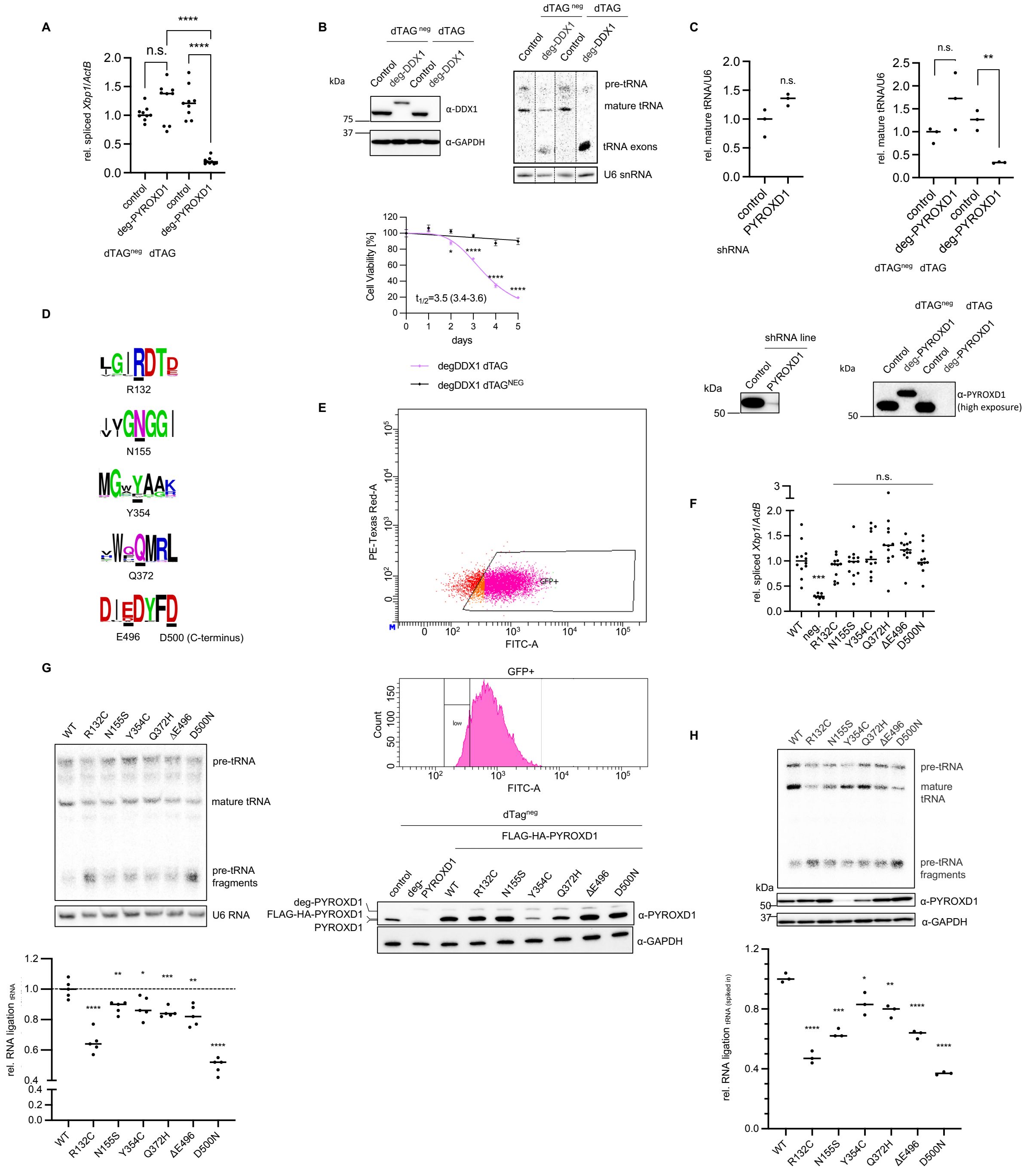


**Figure S2 (related to Figure 2). A**. PYROXD1 was endogenously tagged with FKBP12$^{F36V}$ (here: deg-) tag in HeLa cells, and depleted from cells for 3 days upon addition of degradation (dTAG$^{V-1}$, here: dTAG) drug. Inactive degrader mimic (dTAG$^{V-1_neg}$, here: dTAG$^{neg}$) was used as a negative control. *XBP1* and *Actin* mRNA levels were estimated using RT-qPCR and quantified relative to control cell line treated with dTAG$^{neg}$. Experiment was conducted 9 times. **** p<0.0001. **B**. DDX1 was endogenously tagged and cells were treated as in (A). DDX1 levels were estimated by western blotting, pre-tRNA fragment ligation was followed by northern blotting in RNA extracts. Experiment was conducted 3 times and the representative image displayed. Cell viability was estimated using AquaBluer (N=5). **** p<0.0001. **C**. Quantifications in this figure are based on experiments presented in Figs. S1A and 2B. Briefly, PYROXD1 was silenced for 7 days using shRNA (top left) or degraded for 3 days with dTag drug (top right) and pre-tRNA fragment ligation was monitored by northern blotting of Tyr$^{GTA}$. Mature tRNA/U6 ratio was estimated and quantified relative to designated control lines. ** p<0.01 (student t-test). High exposure western blots reveal traces of PYROXD1 in shRNA line (bottom left), absent from deg-PYROXD1 line (bottom right). **D**. Conservation of amino acids in PYROXD1 across animals: sequence logos display conservation of each residue (underlined) affected in single amino acid patient PYROXD1 variants. **E**. Cells endogenously tagged as in (A) were virally transduced to generate lines constitutively expressing FLAG-HA-PYROXD1 in a manner coupled with GFP. Cells were sorted twice, and each time the population with the lowest fluorescence (orange, "low") was selected for propagation (top and middle – second round of sorting is displayed). Levels of PYROXD1 in the generated lines were estimated via western blotting (bottom). **F**. Cells constitutively expressing FLAG-HA-PYROXD1 variants generated in (E) were depleted of the endogenous PYROXD1 using dTag drug for 3 days, and treated with tunicamycin for 3.5 h. *XBP1* and *Actin* mRNA levels were determined in RNA extracts using RT qPCR, and displayed relative to WT cells. *** p<0.001 (t-test). neg: cells expressing no PYROXD1 construct on top of depletion of endogenous protein. **G**. Cells treated as in (F) were harvested and endogenous pre-tRNA ligation was assessed by northern blotting of RNA extracts, and quantified relative to WT (right). **** p<0.0001 *** p<0.001 ** p<0.01 * p<0.05 (student t-test). Cells generated in (E) were depleted of the endogenous PYROXD1 using dTag drug for 3 days, and northern blotting against $Tyr^{GTA}$ was performed on the RNA extracts as in (B). **** p<0.0001 *** p<0.001 ** p<0.01 * p<0.05 (student t-test). **H**. Cells were treated as in (G), and splicing and ligation of radiolabelled tRNAs spiked into the protein extracts was monitored by urea PAGE and autoradiography, and quantified relative to WT (bottom). Protein amounts are assessed by western blotting. pre-tRNA fragments generated in this assay are distinct from those in Northern blots due to the lack 5'leader and 3'trailer sequences. Experiment was conducted 3 times. **** p<0.0001 *** p<0.001 ** p<0.01 (student t-test)

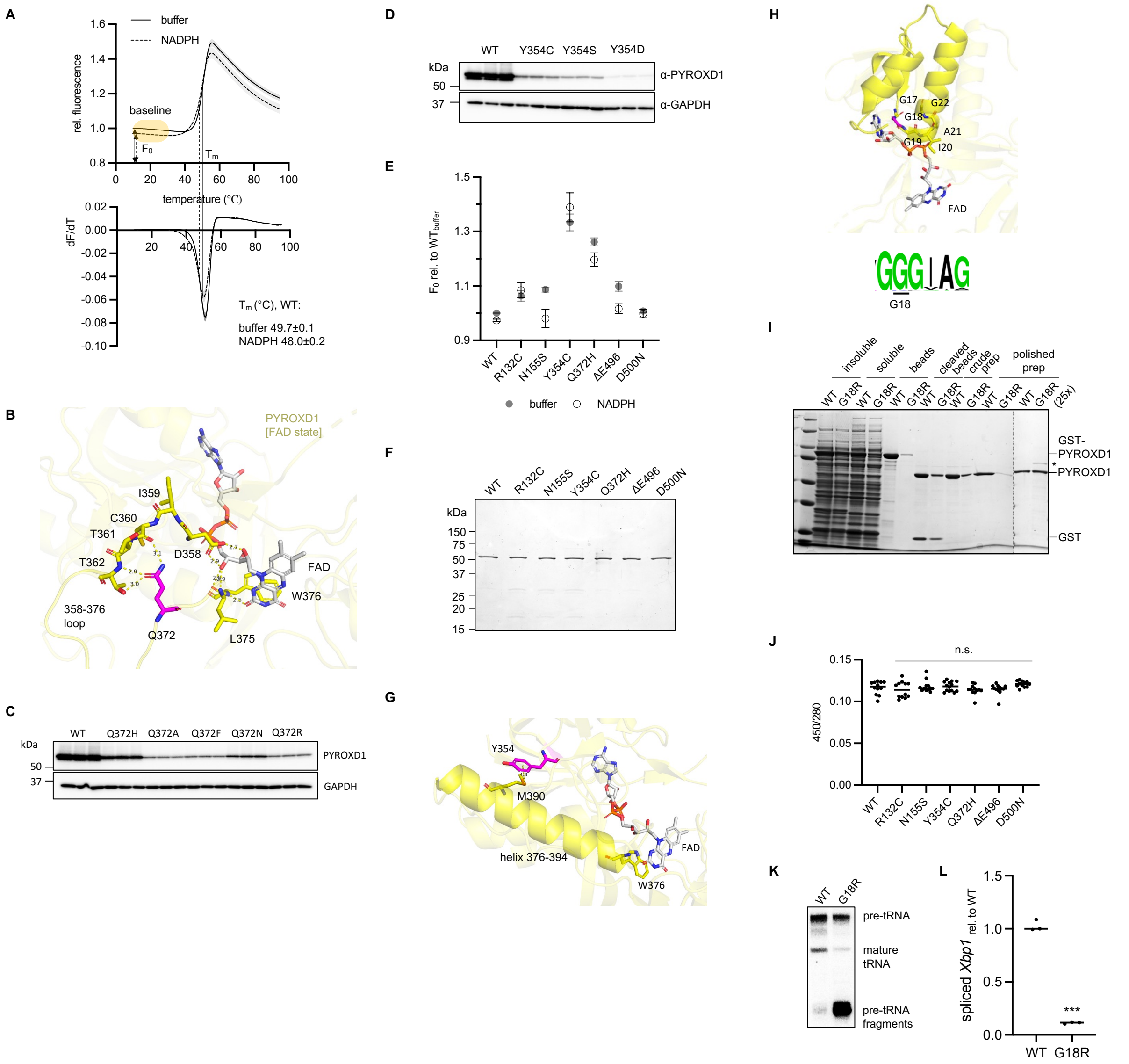


**Figure S3 (related to Figure 3). A.** Recombinant PYROXD1 was expressed in *E. coli* from pGex-3p1 plasmid and purified using GST-affinity capturing, tag cleavage and size exclusion. The protein was incubated with 1 mM NADPH or buffer for 1 h at room temperature as designated. Protein unfolding was followed by SYPRO Orange thermal shift assay. N=14. Each measurement is conducted with a separate aliquot from at least two independent preparations. To account for peak asymmetry, centroid of the derivative curve (bottom) was used to determine the melting point, and average value between 10 and 30°C was used for the baseline (top, see Figs. 3C and S3E). **B**. Q372 is positioned within the 358-376 loop that binds FAD on both ends. Previously published structure (PDB: 6ZK7) was used. **C**. HeLa cells, with PYROXD1 endogenously tagged by $FKBP12^{F36V}$ tag were depleted of endogenous PYROXD1 for 3 days after being transiently transfected with Lipofectamine 3000 with plasmid expressing FLAG-HA-PYROXD1 constructs of interest. Protein levels were assessed by western blotting. **D**. FLAG-HA-PYROXD1 constructs were transiently transfected and analysed as in (C). **E**. Recombinant PYROXD1 variants were expressed, purified and analysed as in (A). Pre-unfolding baseline range of non-derivatised thermal shift curves is displayed. N≥9. Statistical significance. Buffer: R132C * N155S **** Y354C **** Q372H **** ΔE496 ** D500N n.s., NADPH: R132C ** N155S n.s. Y354C **** Q372H **** ΔE496 * D500N n.s. (student t-test) **F**. Polished preparations of recombinant PYROXD1 variants obtained as in (A) were analysed by SDS PAGE followed by InstantBlue staining. **G**. Y354 interplays with 376-394 helix, which starts with FAD-stacking W376 residue. Our previously published structure (PDB: 6ZK7) was used. **H**. G18 resides within Rossmann motif (top) and is largely conserved in animals (bottom). Our previously published structure (PDB: 6ZK7) was used. **I**. GST-PYROXD1 constructs were expressed in *E. coli* and purified by affinity capturing, prescission protease cleavage and size exclusion as in (A). Purification stages were monitored by SDS PAGE followed by InstantBlue staining. **J**. UV-visible spectra of recombinant PYROXD1 proteins were recorded and 450/280 ratio was determined to estimate FAD occupancy. N=14. Each measurement is conducted with a separate aliquot from at least two independent preparations. Student t-test was used for analysis. **K**. PYROXD1 was endogenously tagged with FKBP12$^{F36V}$ tag in HeLa cells, and FLAG-HA-G18R or WT PYROXD1 were stably integrated, resulting in constitutive expression. Endogenous PYROXD1 was depleted for 3 days with dTAG drug and pre-tRNA fragment ligation to mature tRNAs was assessed by northern blotting against Tyr$^{GTA}$. Experiment was conducted 3 times and the representative image displayed. **L**. Cells were treated as in (K), followed by treatment with tunicamycin for 3.5 h. *XBP1* and *Actin* mRNA levels were assessed by RT qPCR in RNA extracts, and the results were expressed relative to WT. N=3. *** p<0.001

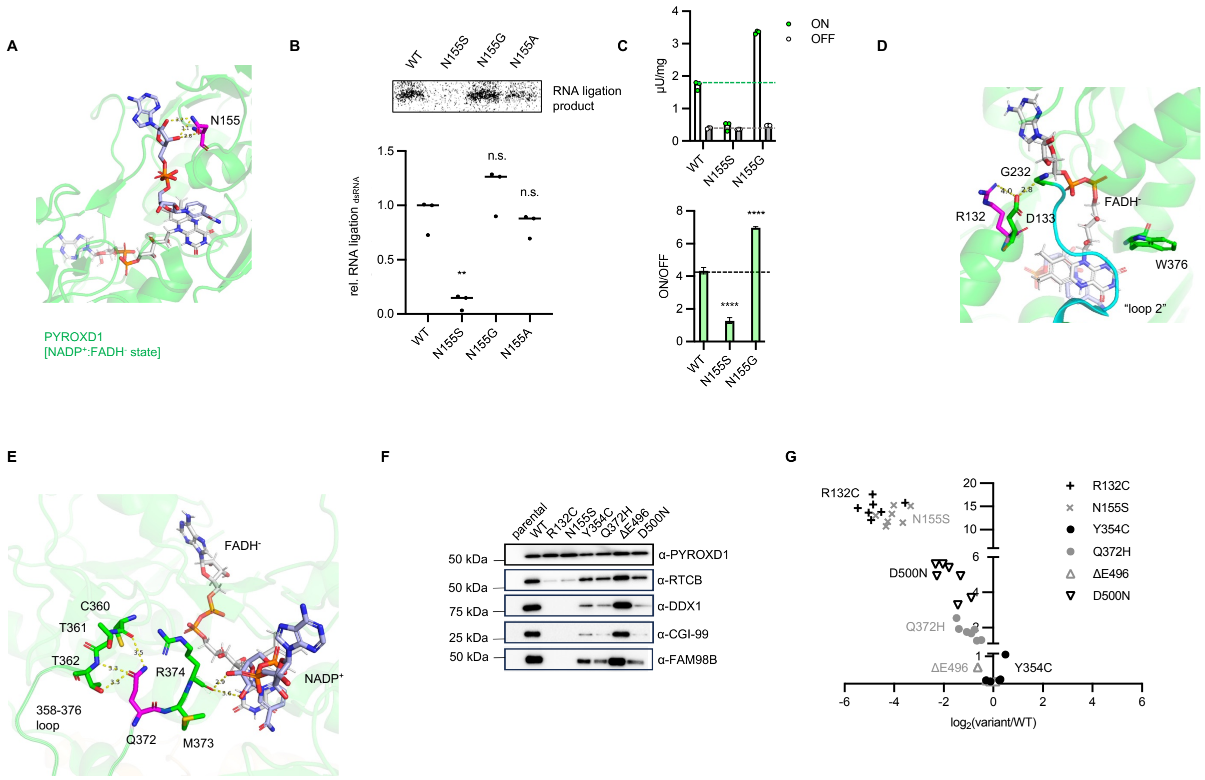


**Figure S4 (related to Figure 4). A.** N155 residue is involved in positioning of NADPH in the active site of PYROXD1. Image is based on our previously published structure (PDB: 8ORJ) **B**. HeLa cells, with PYROXD1 endogenously tagged by FKBP12$^{F36V}$ tag were depleted of endogenous PYROXD1 for 3 days after being transiently transfected with Lipofectamine 3000 with plasmid expressing FLAG-HA-PYROXD1 constructs of interest. dsRNA oligo ligation was conducted in cell extracts, and the products were separated with urea PAGE and visualised by autoradiography. Protein amounts were analysed by northern blotting. N=3. ** p<0.01 (student t-test). The experiment was conducted independently from the one shown in Fig. 4C. **C**. Recombinant PYROXD1 variants were expressed in *E. coli* from pGex-3p1 plasmid and purified from using GST-affinity capturing, tag cleavage and size exclusion. Proteins were incubated with NADPH together with either artificial quinoid substrate menadione or buffer ($O_2$ serving as electron acceptor) to assess the rates of on and off reactions during PYROXD1 reaction cycle. Reaction was monitored for 45 minutes and activity was determined from the linear range of the absorbance loss at 340 nm. Activity unit is defined as the amount of enzyme that converts 1 μmol of NADPH per minute and the protein concentration was determined with Bio-Rad Protein Assay. N=3. Statistical significance: ON N155S *** N155G ***, OFF N155S n.s. N155G * (student t-test) (top). ON/OFF ratios were determined and errors and significance were estimated by bootstrap resampling (bottom). **D**. R132 participates in the interaction network that positions the regulatory loop above NADP$^+$:FADH$^-$ charge transfer complex, potentially affecting accessibility to $O_2$. Image is taken from our previously published structure (PDB: 8ORJ) **E**. Similarly to binding of FAD (see Fig. S3B), residues from 358-376 loop containing Q372 residue participate in NADPH binding. (PDB: 8ORJ) **F**. FLAG-HA-PYROXD1 was immunoprecipitated from HEK293 Flp-In cells upon 1 day of doxycycline-induced over-expression, with lysis buffer and wash buffers supplemented with NADPH. Designated immunoprecipitated proteins were visualised by western blotting. Experiment was conducted five times and representative image displayed. **G**. FLAG-HA-PYROXD1 was immunoprecipitated from HEK293 cells as described in (F) and analysed by mass spectrometry. Each dot represents an average value for an individual subunit of the tRNA-LC across four replicates.

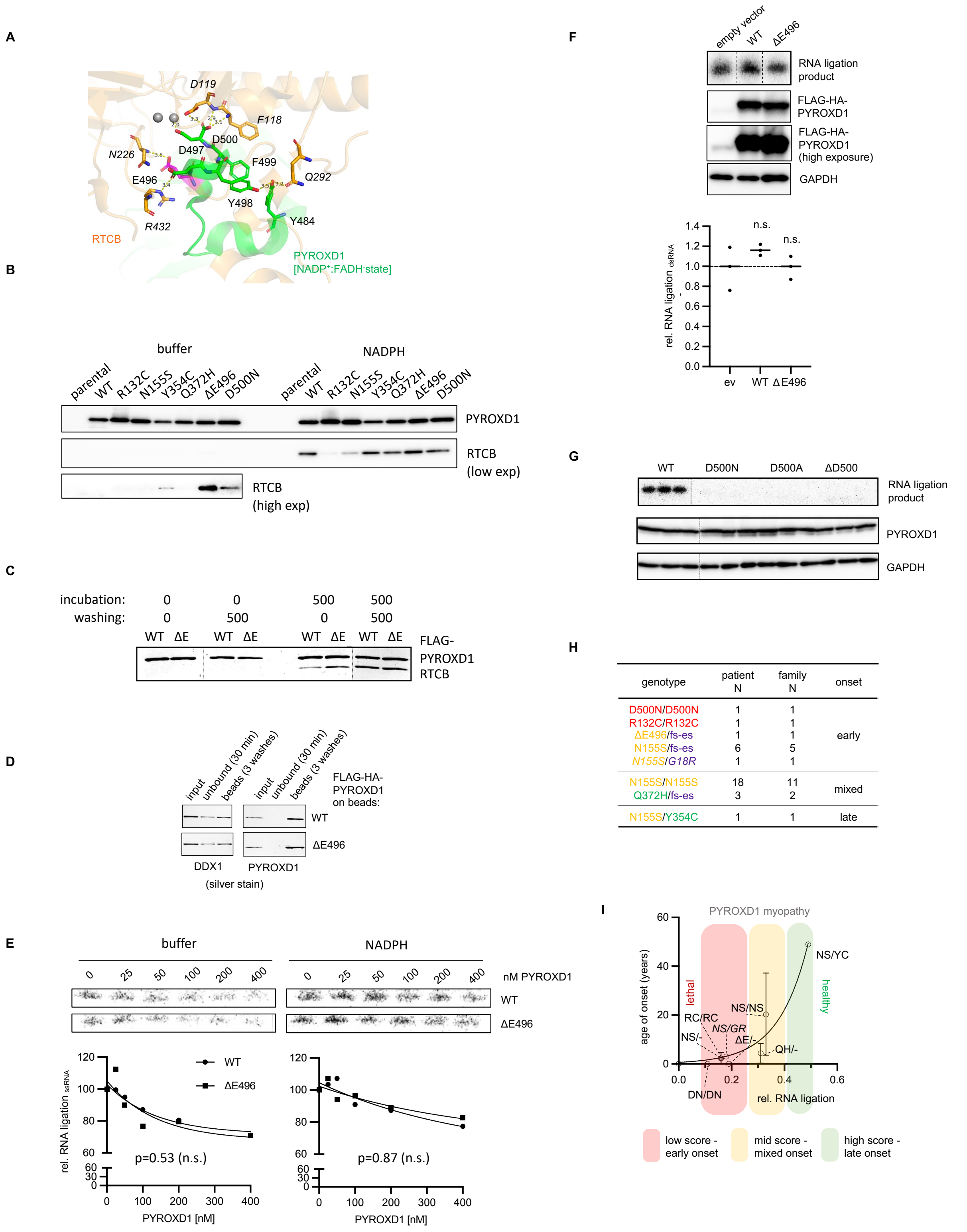


**Figure S5 (related to Fig. 5). A.** C-tail residues interplay with the upstream RBD region for optimal positioning in RTCB binding site, implying potential importance in conformation and flexibility of C-tail for binding kinetics and affinity. Image is derived from our previous structure (PDB: 8ORJ) **B**. FLAG-HA-PYROXD1 was immunoprecipitated from HEK293 Flp-In cells upon 1 day of doxycycline-induced overexpression. Immunoprecipitated proteins were visualised by western blotting. NADPH was included in lysis and washing buffers where designated. Long exposure reveals trace amounts of RTCB in the immunoprecipitates generated without supplementation with NADPH upon prolonged exposure. **C**. Recombinant FLAG-HA-PYROXD1 variants were expressed in *E. coli* from pGex-3p1 plasmid and purified from using GST-affinity capturing, tag cleavage and size exclusion. The protein was incubated with the tRNA-LC (4 subunits, untruncated) and immunoprecipitated; NADPH was included in incubation or washing buffers as indicated (0 and 500 indicating NADPH concentration in µM), and the proteins were visualised by silver staining. **D**. The recombinant FLAG-HA-PYROXD1 and the tRNA-LC, expressed and purified as in (C), were incubated with anti-FLAG beads in the presence of NADPH, the beads were washed, and the buffer was exchanged for NADPH-free solution. The figure presents controls (input sample, unbound fraction and the fraction retained on beads after 3 washes) to the panels shown in Fig. 5B (top right). Proteins are visualised by SDS PAGE and silver staining. **E**. Recombinant tRNA-LC was incubated with an increasing concentration of recombinant PYROXD1 and the ligation activity was subsequently measured upon addition of TCEP, ATP, GTP and radiolabelled ssRNA, followed by urea PAGE and autoradiography. RNA ligation was quantified relative to corresponding sample without PYROXD1. Extra sum-of-squares F test was used for the whole curve comparison. **F**. Control HeLa cells were transiently transfected with Lipofectamine 3000 with plasmid expressing FLAG-HA-PYROXD1 constructs of interest. dsRNA oligo ligation was conducted in cell extracts, and the products were separated with urea PAGE and visualised by autoradiography. Protein amounts were analysed by northern blotting. N=3. Student t-test was used for analysis. **G**. HeLa cells, with PYROXD1 endogenously tagged by $FKBP12^{F36V}$ tag were depleted of endogenous PYROXD1 for 3 days after being transiently transfected with Lipofectamine 3000 with plasmid expressing FLAG-HA-PYROXD1 constructs of interest. dsRNA oligo ligation was conducted in cell extracts, and the products were separated with urea PAGE and visualised by autoradiography. Protein amounts were analysed by northern blotting. N=3. **H**. Combinations of alleles observed in the known PYROXD1 patients are grouped based on the severity of RNA ligation failure in the corresponding variant combinations. All patients with two severe variants or one severe and one moderate variant displayed early disease onset. **I**. Schematic presentation of disease onset of the reported patients against the average ligation score of the variants found in the two alleles, calculated under yet to be verified assumption of additive effects, and based on the average values in Figs. 2F and 3G. The exponential fit is used only for guidance and not as a high confidence curve. Error bars represent SD of all known patients with the described genotype. Lethal and healthy outcomes are predictions consistent with the absence of known patients residing in the designated RNA ligation areas (approximately <0.1 and >0.5, respectively). RC: R132C, NS: N155S, YC: Y354C, QH: Q372H, ΔE: ΔE496, DN: D500N, - : frameshift or exon skipping variant; GR: G18R (reported only in a conference abstract).
